# Spatial domain analysis predicts risk of colorectal cancer recurrence and infers associated tumor microenvironment networks

**DOI:** 10.1101/635730

**Authors:** Shikhar Uttam, Andrew M. Stern, Samantha Furman, Filippo Pullara, Daniel Spagnolo, Luong Nguyen, Albert Gough, Christopher J. Sevinsky, Fiona Ginty, D. Lansing Taylor, S. Chakra Chennubhotla

**Author notes:** (Shikhar Uttam); (S. Chakra Chennubhotla).

## Abstract

An unmet clinical need in solid tumor cancers is the ability to harness the intrinsic spatial information in primary tumors that can be exploited to optimize prognostics, diagnostics and therapeutic strategies for precision medicine. We have developed a transformational spatial analytics (SpAn) computational and systems biology platform that predicts clinical outcomes and captures emergent spatial biology that can potentially inform therapeutic strategies. Here we apply SpAn to primary tumor tissue samples from a cohort of 432 chemo-naïve colorectal cancer (CRC) patients iteratively labeled with a highly multiplexed (hyperplexed) panel of fifty-five fluorescently tagged antibodies. SpAn predicted the 5-year risk of CRC recurrence with a mean area under the ROC curve of 88.5% (SE of 0.1%), significantly better than current state-of-the-art methods. SpAn also inferred the emergent network biology of the tumor spatial domains revealing a synergistic role of known features from CRC consensus molecular subtypes that will enhance precision medicine.

## Introduction

Colorectal Cancer (CRC) is the second most common type of cancer and the third leading cause of cancer-related deaths worldwide.^1^ This multi-factorial disease like other carcinomas, develops and progresses through the selection of epithelial clones with the potential to confer malignant phenotypes in the context of a reciprocally coevolving tumor microenvironment (TME) comprising immune and stromal cells.^2–4^ CRC patients are staged using the well-established tumor-node-metastases (TNM) classification.^5, 6^ However, there is significant variability in patient outcomes within each stage. For example, CRC will recur in up to 30% of Stage II patients despite complete tumor resection, no residual tumor burden and no signs of metastasis.^7^ In contrast, more advanced CRC has been known to show stability or indeed even to spontaneously regress.^7, 8^

The intrinsic plasticity of the TME underlying this variability in outcome is controlled by complex network biology emerging from the spatial organization of diverse cell types within the TME and their heterogeneous states of activation.^3, 9–11^ The important role of the TME in CRC progression and recurrence has recently been highlighted by the identification of four consensus molecular subtypes (CMS) ^12, 13^, functional studies defining the critical role of stromal cells in determining overall survival,^14^ and the development of Immunoscore®^14^ which quantifies tumor-infiltrating T-lymphocytes in different regions of the tumor and associates their infiltration with CRC recurrence.^15, 16^ However the TME can be further harnessed to significantly improve CRC prognosis through the identification of biomarkers mechanistically linked to disease progression and the development of novel therapeutic strategies.

Deeper understanding of the TME may arise from imaging methods capable of labeling > 7 cellular and tissue components in the same sample (hyperplexed^17^ (HxIF) fluorescence and other imaging modalities).^17–21^ To fully extract the intrinsic information within each primary tumor we have developed a <u>sp</u>atial <u>an</u>alytics (SpAn) computational and systems pathology platform applicable to all solid tumors to analyze the spatial relationships throughout TME signaling networks. SpAn constructs a computationally unbiased and clinical outcome-guided statistical model enriched for a subset of TME signaling networks that are naturally selected as dependencies of the corresponding malignant phenotype.^22–25^ Traditionally, advances in experimental and systems biology have been made by identifying associations between differential biomarkers expressions/correlations, or clusters in T-SNE plots, with particular outcome-specific malignant phenotypes, such as cancer progression or recurrence phenotypes. Instead of using this association-driven paradigm, SpAn introduces an outcome-driven approach to predict 5-year risk of CRC recurrence in patients with resected primary tumor that also enables inference of recurrence-specific network biology.

## Results

### Hyperplexed immunofluorescence imaging of tissue microarrays generates multidimensional spatial proteomics data with sub-cellular resolution

The acquired data were generated using GE Cell DIVE^TM^, also known as MultiOmyx^20^, (GE Healthcare, Issaquah, WA) hyperplexed^17^ immunofluorescence (HxIF) imaging and image processing workflow instrument. As previously described,^20^ Cell DIVE^TM^ can perform hyperplexed imaging of greater than 60 biomarkers via sequentially multiplexed imaging of 2 to 3 biomarkers plus DAPI nuclear counterstain through iterative cycles of label–image–dye-inactivation visualized in Supplementary Fig. S1.^20^ (See Methods for more details.) Extensive validation of this approach has demonstrated that a majority of epitopes tested are extremely robust to the dye inactivation process. The biological integrity of the samples were preserved for at least 50 iterative cycles.^20^

In this study we use 55 biomarkers, which include markers for epithelial, immune and stromal cell lineage, along with those in categories, which include 1. biomarkers sampling the network biology of signalling pathways, 2. biomarkers associated with extracellular transport and metabolism, 3. biomarkers associated with tumour suppressive potential, 4. biomarkers associated with oncogenic potential, 5. biomarkers associated with cell-cell adhesion, cellular and stromal structure, 6. biomarkers associated with post-translational modifications (PTM), and 7. biomarkers associated with cell types and their states. They are detailed in Fig. 1, with additional details in Supplementary Table S1. Figure 2a shows the HxIF image stack of a 5 µm thick and 0.6mm wide tissue microarray^26^ (TMA) spot from resected primary tumor of a Stage II CRC patient labelled with the 55 biomarkers plus DAPI. Figure 2b highlights a sub-region of this patient TMA spot enabling optimal visualization of the 55 HxIF biomarker images resulting from the iterative label– image–chemical-inactivation cycles.

**Figure 1:**
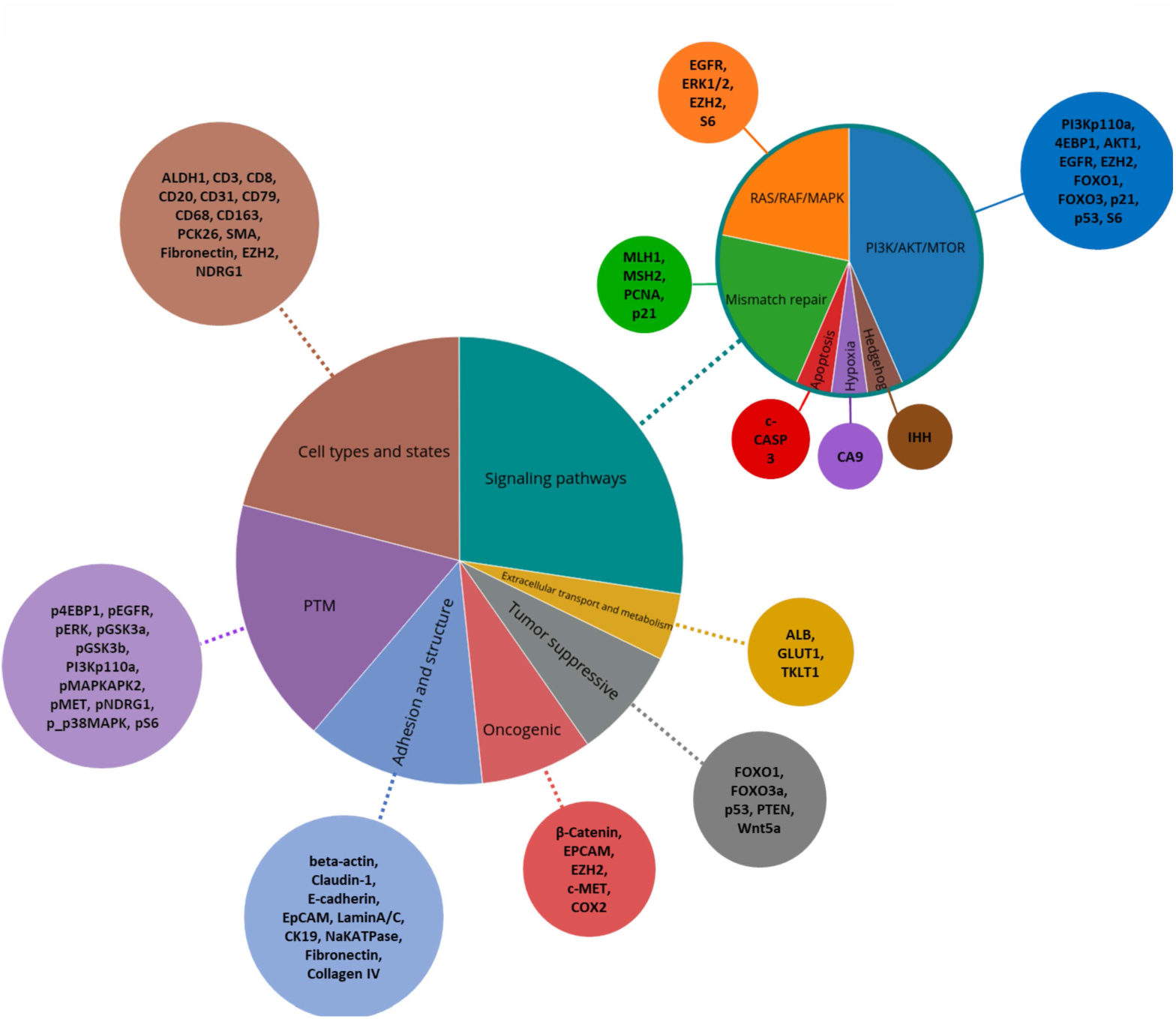
Biomarker selection and organization. Fifty-five biomarkers along with DAPI were imaged using the Cell DIVE platform.^20^ The biomarkers were selected from seven broad categories that include 1. biomarkers sampling the network biology of signaling pathways (PI3K/AKT/MTOR^s1^, RAF/RAF/MEK^s2^, Mismatch Repair^s3^ (MMR), Hedgehog signaling^s4^, Hypoxia-signaling^s5^, Apoptosis^s6^); 2. biomarkers associated with extracellular transport and metabolism (Albumin^s7-s9^, GLUT1^s10,s11^, TKLP1^s12,s13^); 3. biomarkers associated with tumor suppressive potential (FOXO1^s14,s15^, FOX03^s15,s16^, p53^s17^, PTEN^s18^, Wnt5a^s19^); 4. biomarkers associated with oncogenic potential (EPCAM^s20^, COX2^s21,s22^, c-MET^s23,s24^, Beta-Catenin^s24,s25^, EZH2^s26-s30^); 5. biomarkers associated with cell-cell adhesion, cellular and stromal structure (Beta-actin^s31^, Claudin-1^s32^, E-cadherin^s33^, EPCAM^s20^, Lamin A/C^s34,s35^, CK19^s36^, NaKATPase^s37^, Fibronectin^s38^, Collagen IV^s39^); 6. biomarkers associated with post-translational modifications (PTM)^s40^ (pE4BP1, pMET, pERK1/2, pMAPKAPK2, p38MAPK, pEFGR, pGSK3a/b, pNDRG1, pS6); and 7. biomarkers associated with cell types and their states (ALDH1^s41,s42^, CD20^s43,s44^, CD68^s45^, CD163^s45^, CD31^s46^, CD79^s47^, EZH2^s26–s30^, CD3^s48^, CD8^s49,s50^, PCK26^s51^, SMA^s52,s53^, Fibronectin^s38^, NDRG1^s54^). The references are detailed in Supplementary Table S1 and are indicated here with a prefix ‘s’.

**Figure 2:**
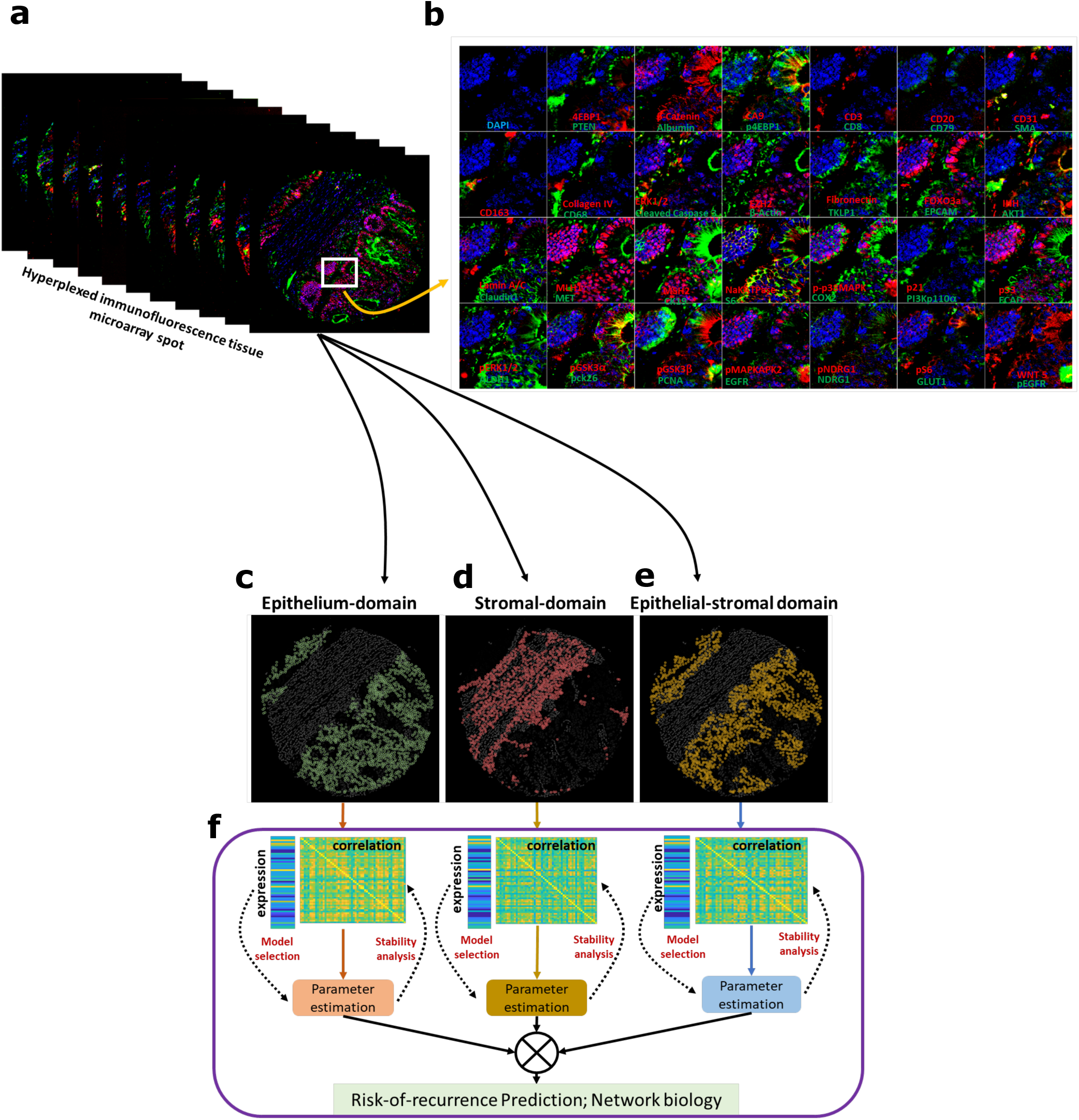
Spatial analytics (SpAn) platform based on hyperplexed immunofluorescence imaging for predicting risk of 5-year CRC recurrence. (a) Hyperplexed image-stack of a TMA spot generated by iteratively multiplexed (Fig. S1) HxIF imaging using the Cell DIVE platform.^20^ (b) Close-up view of a TMA region in (a) outlined in white (approx. 110µm by 110µm), labelled with 55 biomarkers (plus DAPI nuclear counterstain) that include epithelial, immune and stromal cell lineage, subcellular compartments, oncogenes, tumour suppressors, and posttranslational protein modifications described in detail in supplementary Table S1. Hyperplexed imaging is implemented via iterative label–image–dye-inactivation immunofluorescence cycle (see Methods and Fig. S1). (c-e) Dissection of the TMA spot into three spatial-domains (epithelial, stromal, and epithelial-stromal domains) identified and segmented using structural biomarkers (see Methods and Fig. S2). (f) For each of the three spatial domains both expressions of the 55 biomarkers and their Kendall rank-correlations both within and across the cells together defined the domain-specific features. L1-norm based penalized Cox regression was used for model selection, while L2-penalty was used for final model parameter (coefficients) estimation. The stability of the model was tested at the 90% concordance level, and the parameters were reevaluated for final construction of the SpAn spatial model.

Cell DIVE^TM^ was employed to generate HxIF image stacks of FFPE tissue microarrays from resected tissue samples from 432 chemo-naïve CRC patients at single cell resolution. This 55-dimensional spatial profiling of the patient-level tumor microenvironment served as input in our study. The 432 retrospectively acquired chemo-naïve CRC patient cohort included patients in Stage I through Stage III of CRC primary tumor growth between the years of 1993 and 2002 acquired from Clearview Cancer Institute of Huntsville Alabama. As shown in Supplementary Table S2, the median patient age and gender proportions were similar across all stages, with CRC recurring in 65 patients. The outcome distribution of the patients and their clinical attributes across the CRC stages are detailed in Table S2. The use of chemo-naïve (no administration of neoadjuvant or adjuvant therapies for the 5+ years of follow-up) CRC patient cohort provides SpAn the opportunity to interrogate unperturbed primary tumor biology.

### SpAn uses HxIF spatial proteomics data to learn recurrence-guided and spatially informed prognosis of CRC recurrence

SpAn performs a virtual three-level spatial-dissection of the tumor microenvironment, by first explicitly decomposing the TMA into epithelial and stromal regions as detailed in Methods and illustrated in Fig. S2. The cells in the epithelial region are identified using E-cadherin cell-cell adhesion labeling and pan-cytokeratin, with individual epithelial cells segmented using a Na^+^K^+^ATPase cell membrane marker, ribosomal protein S6 cytoplasmic marker, and DAPI-based nuclear staining. The resulting epithelial spatial domain of the TMA in Fig. 1a is shown in Fig. 2c. The remaining cells are assigned to the stromal domain and are visualized in Fig. 2d. These stromal cells have diverse morphologies.^20^ Based on the epithelial and stromal domains, SpAn also identifies a third epithelial-stromal domain, shown in Fig. 2e, to explicitly capture a 100 µm boundary wherein the stroma and malignant epithelial cells interact in close proximity. Together these three intra-tumor spatial domains comprise the virtual three-level spatial dissection of the tumor microenvironment that forms the basis for the SpAn spatial model overviewed in Fig. 2f and detailed in Fig. 3.

**Figure 3:**
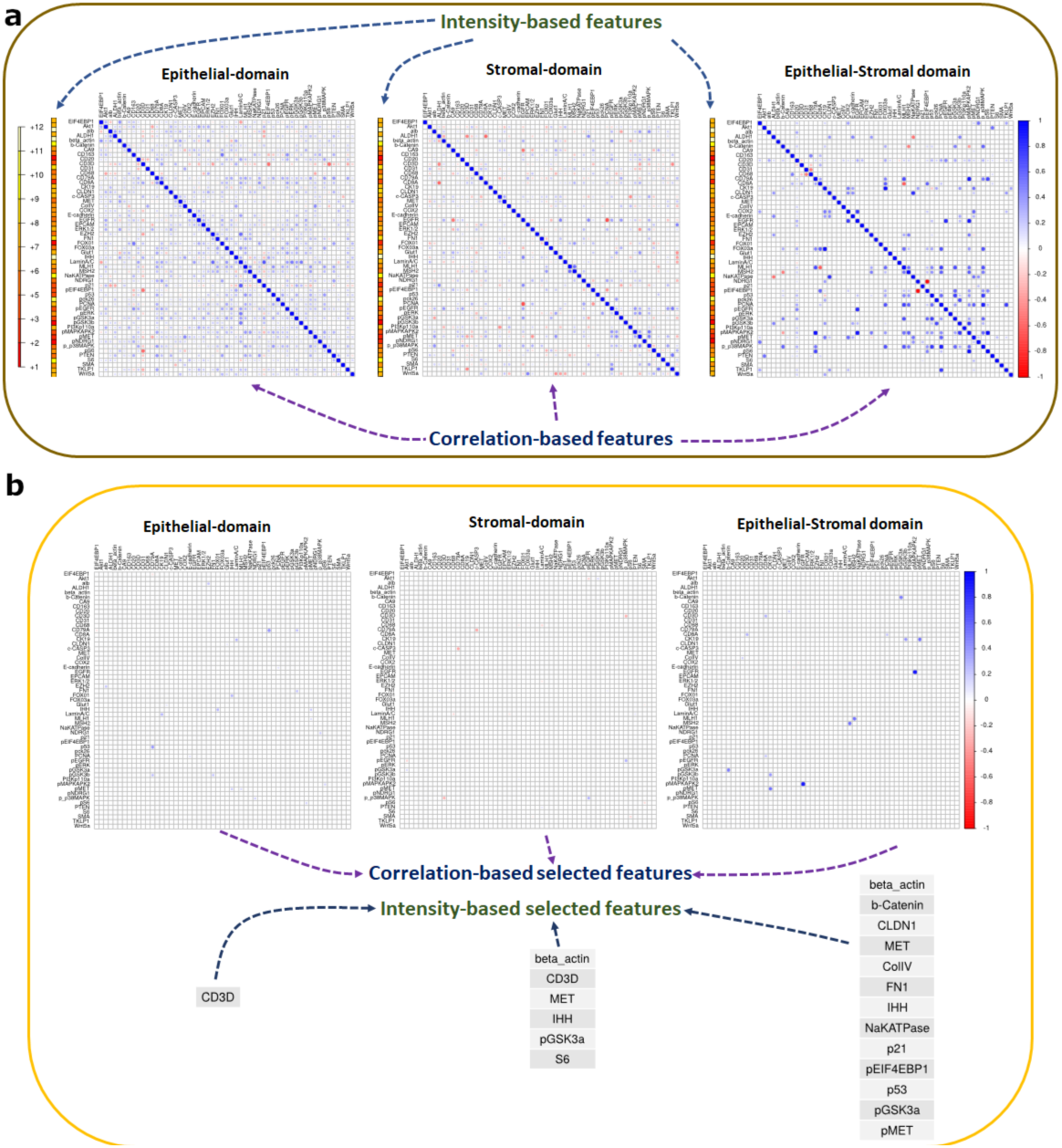
SpAn domain-specific feature selection. (a) Individual mean protein-expression intensity profile depicted as a vector and pairwise Kendall rank-correlations between protein expressions – visualized as a matrix for each of the three spatial domains. The protein expressions are shown in log scale. To prevent inclusion of false-positive protein expression, only intensities above the 85^th^ percentile were considered as expressions and used to compute the correlations. (See Methods.) (b) Features, including both expressions and correlations, selected by SpAn based on L1-penalized Cox regression used for model selection. The selected features were consistently concordant at the 90% level with the recurrence outcome.

Utilizing expression of the 55 hyperplexed biomarkers, SpAn first computes the corresponding 55 mean intensities and 1485 Kendall rank-correlations as features characterizing each of the three spatial domains (see Fig. 3a). The mean intensity captures the average domain-specific expression profile of each biomarker, while the Kendall rank-correlations^27^ measure strength of association between any two biomarkers without presuming linearity. (See Method for details.) Importantly, computation of domain-specific rank-correlations as explicit features for SpAn is used in place of the more typical approach of implicitly incorporating correlations as interactions between covariates (average biomarker expressions) within the prediction model^28^. These explicit features not only detect the association between two biomarkers presumably mediated by intracellular and intercellular networks all within the same spatial domain but also by mediators (e.g., exosomes) derived from another spatial domain. As an example, SpAn finds enrichment of KEGG ‘microRNAs in cancer’ pathway in the epithelial and epithelial-stromal domains (Figure 5), while concurrently selecting correlation between CD163 and PTEN as a feature in the stromal domain for recurrence prognosis (see Fig. 4). As has been reported in gastrointestinal cancers, tumor cell derived exosomal miRNAs mediate crosstalk between tumor cells and the stromal microenvironment, and induce polarization of the macrophages to the anti-inflammatory and tumor-supportive M2 state via activation of the PTEN-PI3K signaling cascade under hypoxic conditions resulting in enhanced metastatic capacity.^29, 30^

**Figure 4:**
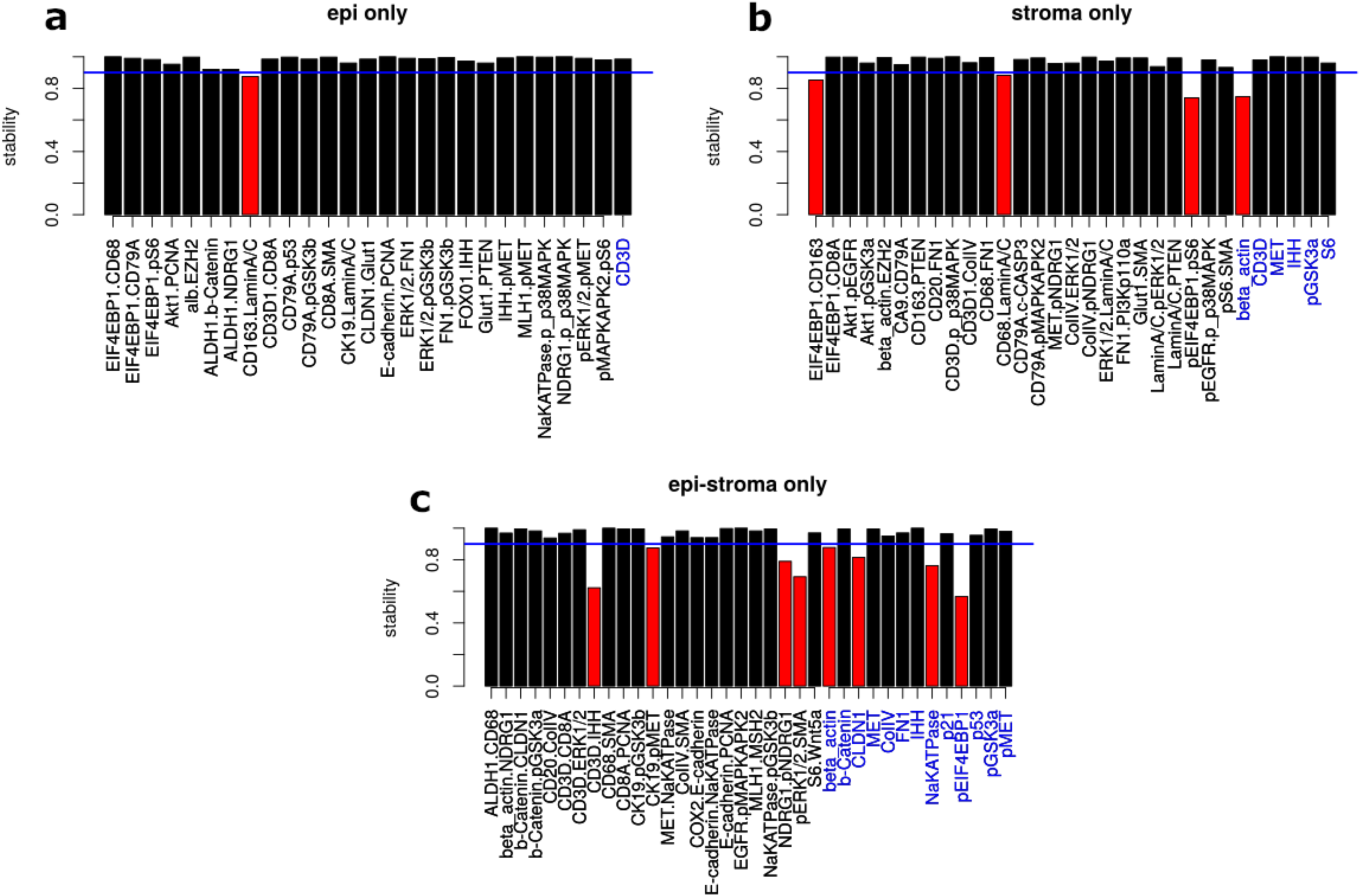
SpAn stability analysis. Stability analysis of the selected features for each of the spatial domains, with only those features from the ones selected in Fig. 3b that maintain their sign in 90% of the 500 bootstrap runs are included as input into the SpAn spatial model. These features are visualized in black in the three bar graphs. Features selected in Fig. 3b that did not meet this criterion are shown in red.

**Figure 5.**
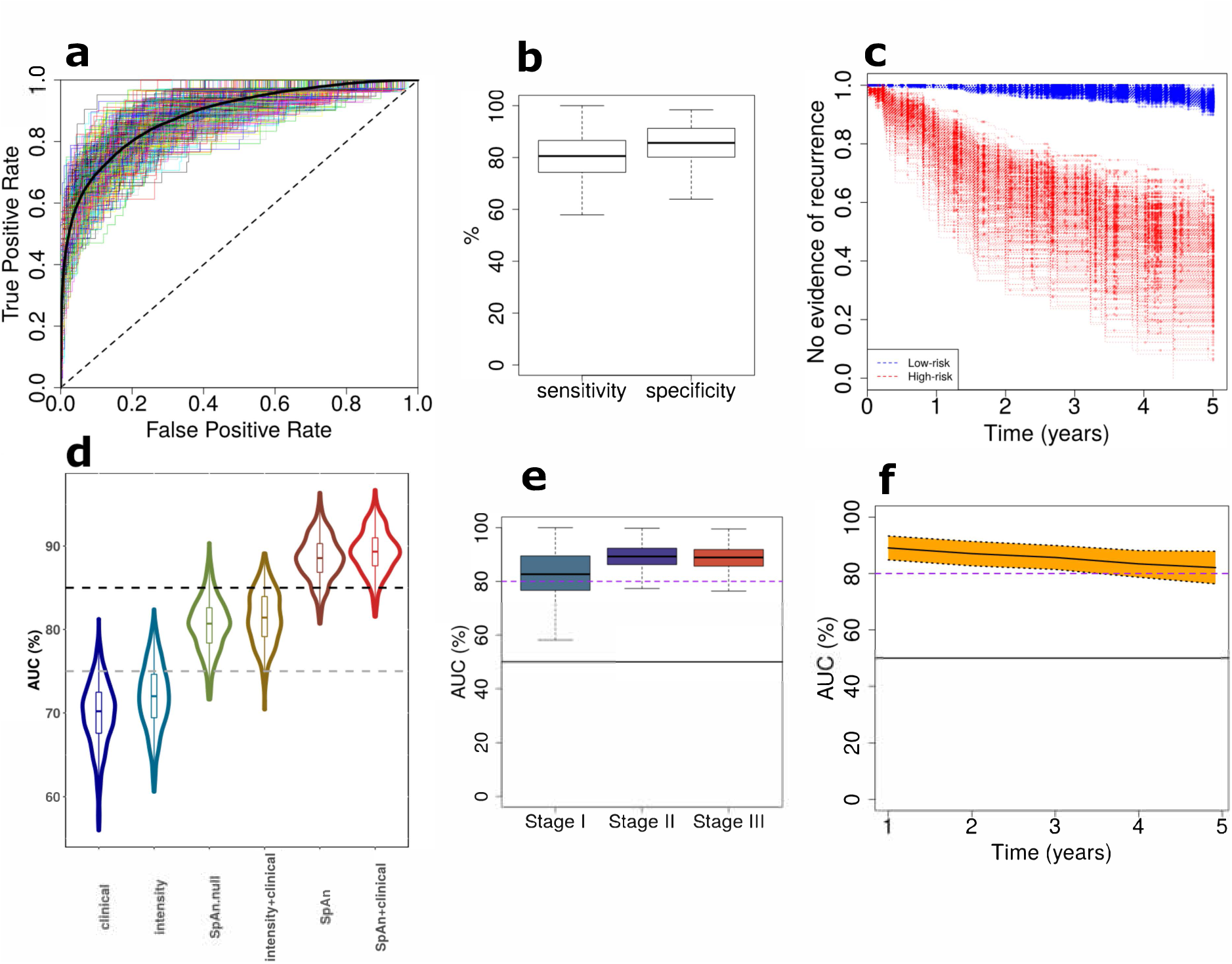
Performance of SpAn platform for predicting risk of 5-year CRC recurrence. (a) SpAn receiver operating characteristics (ROC) curves for predicting risk of 5-year CRC recurrence in patients with resected CRC primary tumor. The plot shows ROC curves for 500 bootstrap runs with independent training and validation sets. Mean area under the ROC curve is 88.5% with a standard error of 0.1%. (b) Boxplots of the sensitivity and specificity values for the same 500 bootstrap runs as in (a) with the operating point on the ROCs chosen by minimizing overall misdiagnosis rate. The mean sensitivity and specificity values respectively are 80.3% (standard error of 0.4%) and 85.1% (standard error of 0.3%). (c) Kaplan-Meier recurrence-free survival curves for each of the 500 bootstrap runs for patients identified by SpAn at low and high-risk of five-year CRC recurrence. (d) AUC violin and box plots of the bootstrapped ROC curves achieved by six CRC recurrence prediction models. The models are 1. clinical model, 2. biomarker expression model (denoted by intensity), 3. SpAn.null model (denoting SpAn without spatial-domain context), 4. biomarker expression + clinical model, 5. SpAn, and 6. SpAn + clinical model. 500 bootstraps were used. Fig. S7 lists the statistical significance of the pairwise performance comparisons between all the six models. The gray and black dashed lines represent ROC curves with AUC of 75% and 85% respectively. (e) Boxplots of stage-based area under the 500 bootstrapped ROC curves showing the stable stage-based clinical performance of SpAn. (f) Stable temporal performance of SpAn illustrated by the time-dependent AUC values plotted as a function of time in years. The 95% confidence interval computed using the 500 bootstraps is also shown by the yellow shaded area around the mean time-dependent AUC values depicted by the solid black line.

SpAn then uses CRC recurrence-guided learning to determine those specific spatial domain features that constitute the optimal subset for prognosis via model selection based on L1-penalized Cox proportional hazard regression method (Figure 3b).^31, 32^ See Methods for details on penalized regression, Fig. S3 for validity of the proportional hazard assumptions, and Figure S4 for determination of threshold for concordance with recurrence outcome. A follow-up analysis of the selected features is performed to test the stability of their contribution to recurrence prognosis through testing the stability of the sign of the corresponding coefficients at the 90% threshold. The final domain-specific features are shown in Fig. 4 and supplementary Table S3 with additional details described in Methods.

The coefficients that control the contribution of the selected features to each of the domain-specific models for assessing recurrence outcome were learned under L1 penalization and their values are, therefore, dependent on all 1540 features. To remove this dependence, SpAn relearns each of the three domain-specific model coefficients using L2 penalty in our penalized Cox regression model with only the optimally selected features as input. This L2-regularized learning allows SpAn to estimate optimal contribution of the selected features that are 90% concordant with the recurrence outcome. The resulting domain-specific coefficients are shown in Supplementary Fig. S5. As detailed in Methods, SpAn combines these domain-specific features weighted by their corresponding coefficients into a single recurrence guided spatial domain prognostic model, whose performance is shown in Fig. 5a. The results were obtained by bootstrapping (sampling with replacement) patient data set to generate 500 pairs of independent training and testing sets using stratified sampling that ensured the proportion of patients in whom cancer recurred in each of the five years remained the same in each bootstrap. For each bootstrap, SpAn used the training data for learning and the independent testing data to compute the receiver operating characteristic (ROC) curve. These ROC curves are shown in Fig. 5a along with the mean ROC curve. The mean area under the curve (AUC) for bootstrapped ROC curves is 88.5% with a standard error of 0.1%, demonstrating the stable performance of SpAn. We also maximized Youden’s index^33^ to identify the clinically relevant operating point on the ROC curves that minimized the overall misdiagnosis rate. Figure 5b shows the resulting sensitivity and specificity values for all bootstrap runs, with mean values respectively of 80.3% (standard error of 0.4%) and 85.1% (standard error of 0.3%). High specificity limits SpAn from misidentifying no-evidence-of-disease patients as being at high risk of CRC recurrence, while at the same time good sensitivity allows SpAn to not miss high-risk patients. This is emphasized by a high positive likelihood ratio value of 7.2 (standard error of 0.23), which quantifies the large factor by which odds of CRC recurring in a patient go up, when SpAn identifies the patient as being at risk of CRC recurrence. At the same time a small negative likelihood value of 0.22 (standard error of 0.003) quantifies the decrease in odds of CRC recurrence in a patient when SpAn identifies the patient as being at low-risk. Finally, these results are brought together in Fig. 5c, which show the large separation in recurrence-free survival curves of patients identified by SpAn at low- and high-risk of five-year CRC recurrence.

### Validating the rationale behind SpAn

The rationale behind our ‘virtual-dissection followed by combination of the three specific spatial domains’ approach is motivated by the acknowledged active role of the microenvironment and its spatial organization, and the differential role played by the epithelial and stromal domains in tumor growth and recurrence.^2, 3, 14, 34^ We tested the validity of this rationale within the context of our data by comparing the performance of SpAn with the null model, which is based on recurrence-guided learning of the spatially undissected patient TMA spot. We note that the learning procedure for the null model was identical to the domain-specific learning within SpAn. Additionally, we also compared the performance of SpAn with four other models that included a clinical model, biomarker expression model, biomarker expression + clinical model, and SpAn + clinical model. The input into the clinical model were clinical features associated with age, gender and TNM stage, and learning procedure was based on Cox proportional hazard regression.^35^ The biomarker expression model input were biomarker expression intensities alone and the learning procedure was identical to SpAn. The biomarker expression + clinical, and SpAn + clinical models respectively combined biomarker expression and SpAn with the clinical model. Figure 5d shows the AUC violin and box plots of the bootstrapped ROC curves achieved by each model. The figure illustrates the improvement SpAn achieves over the performance of other models. To quantify the statistical significance of this improvement, we performed Dunn’s pairwise multiple comparison post-hoc analysis between the models based on non-parametric Kruskal-Wallis test.^36^ Our analysis shows that the improvement in performance achieved by SpAn over all other models is statistically significant at the 99% confidence interval with a p-value much less than 0.005 (Fig. S7). We specifically note that this is true for SpAn performance in comparison to the null model based on the spatially undissected patient TMA spot without spatial-domain context. This improved performance of SpAn highlights the importance of explicitly modeling the epithelial, stromal and epithelial-stromal spatial domains associated with the TME. Interestingly, beyond supporting our rationale, this comparative test also demonstrates that joint utilization of biomarker expressions and their correlations results in superior performance of both SpAn and its null model over clinical features and biomarker expressions alone (also see Fig. S6). We note that published state-of-the-art approaches that include Immunoscore®^15, 16^ rely on biomarker expressions. Finally, we observe that the marginal performance improvement over SpAn achieved by including clinical features with SpAn – the SpAn + clinical model – is not statistically significant with a p-value of 0.082.

### SpAn is highly predictive of 5-year CRC recurrence for TNM Stages I-III in individual patients

The ability to identify patients in whom CRC will recur, especially for those patients in Stages II and III of tumor progression is highly clinically relevant. Figure 5e shows that SpAn can consistently identify patients in whom risk of CRC recurrence is high for Stages I through III, with mean AUC of bootstrapped ROC curves for the three stages respectively being 82.1%, 89.4% and 88.6%. Standard error of these mean AUC values respectively is 0.4%, 0.2%, and 0.2%, demonstrating the stability of SpAn performance. Although the overall performance across all three stages is highly significant with the potential of improving prognosis, the relative reduction in Stage I performance is a consequence of a small cohort of only ten patients in Stage I with CRC recurrence.

The ability of SpAn to predict risk of recurrence in individual patients from all three Stages, is relevant in the context of administering adjuvant therapy, especially for Stage II patients. Current guidelines for treating Stage II CRC patients from The National Comprehensive Cancer Network (NCCN),^37^ the American Society of Clinical Oncology (ASCO),^38^ and the European Society of Medical Oncology (ESMO)^39^ do not recommend routine adjuvant chemotherapy for Stage II patients, but do state that it should be considered for sub-population of Stage II patients that are at higher risk and might benefit from being put on adjuvant therapy regimen.^40^ The personalized prognostic potential of SpAn implies that we could triage Stage II patient cohorts into low and high-risk groups, with the latter being further considered for therapy. Furthermore, SpAn could help with postoperative surveillance of high-risk Stage II patients with more intensive follow-up regimes.^41^

While 20% to 30% of Stage II CRC patients are at high-risk of recurrence, there are Stage III patients that have good prognoses of stable 5-year recurrence-free survival. SpAn, therefore, could also be used to fine-tune their postoperative surveillance and adjuvant chemotherapy regimens.

### Prognostic performance of SpAn remains stable over the 5-year time period

A majority of CRC recurrence occurs in the first five years, with 90% occurring in the first four.^42, 43^ We, therefore, consider the time-dependent performance^44^ of SpAn during the first five-year period. Figure 5f plots the AUC for time-dependent ROC performance. The performance of SpAn in predicting risk of recurrence remains consistent and stable (95% confidence interval shown) with only a small, and gradual reduction in time-dependent AUC values as we move away from the resection and imaging timepoint. This result suggests SpAn captures the critical biological underpinnings of recurrence in the primary tumor. Supplementary Fig. S8 shows the time-dependent AUCs for domain-specific temporal performance of SpAn.

### SpAn identifies spatial domain networks as emergent properties that explain the robust ability to distinguish patients in which CRC recurs

Given the highly prognostic performance of SpAn, we took a systems perspective to understand and to explain the underlying network biology responsible for this performance within each of the three spatial domains. For each domain, we quantified the unique associations between biomarkers included in the selected features through partial correlations between every biomarker pair, when controlling for other biomarkers as described in Methods. This approach was performed for all patients. The resulting partial correlation for every biomarker pair was separated into two groups according to no-evidence-of-CRC and CRC-recurrence patient cohorts and the information distance based on Jensen-Shannon divergence^45^ was computed between them (see Methods for more details). The resulting domain-specific distance matrices, shown in Figs. 6a-c, define associated graphs with the nodes being the biomarkers and edge weights quantifying the differential change, the information distance, in biomarker association between patients in which CRC recurred and those in which there was no evidence of recurrence. The stronger the weights, the larger the distance and the more significant the differential change in association between the two markers for the two patient cohorts. We defined the graphs generated by the distance matrices thresholded at the 99^th^ percentile as the spatial domain networks that were most significant for CRC recurrence prognosis. Figures 6d-f show the resulting networks for the three spatial domains that reveal the heterogeneous nature of the cell populations and signaling pathways leveraged by SpAn in CRC recurrence prognosis.

**Figure 6.**
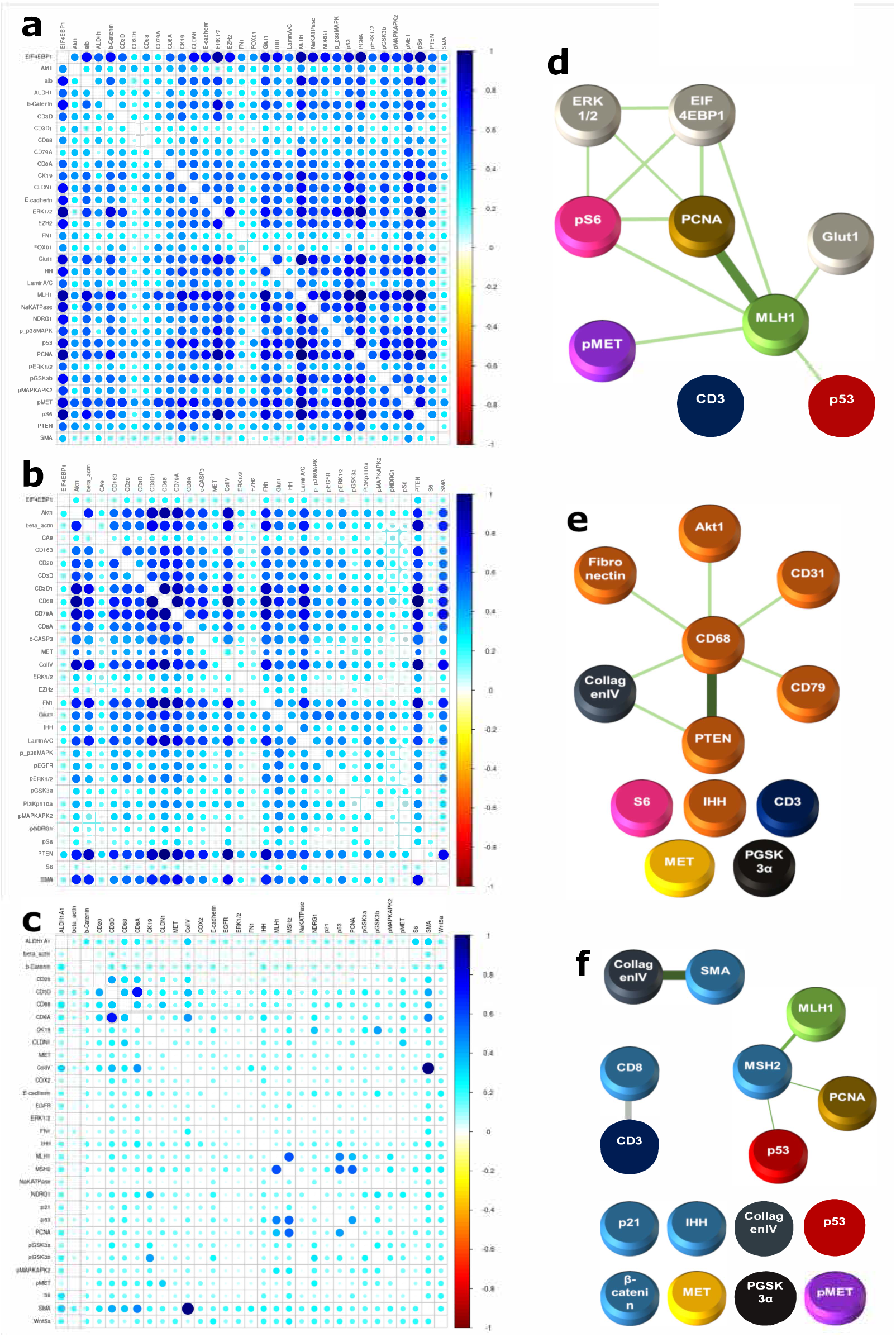
SpAn based computational and systems derived spatial domain networks. (a) Epithelial domain, (b) Stromal domain, and (c) Epithelial-stromal domain Jensen-Shannon divergence matrices that show the information distance of partial correlations (computed for biomarkers selected by recurrence-guided SpAn feature-selection and stability analysis) between patients in the no-evidence-of-CRC and CRC-recurrence cohorts. (d) Epithelial domain, (e) Stromal domain, and (f) Epithelial-stromal domain spatial domain networks obtained by thresholding the corresponding spatial domain information distance matrices at the 99^th^ percentile, which identify differential change most significant for CRC recurrence prognosis. All three spatial-domain networks include disconnected subnetworks with isolated nodes. These nodes correspond to biomarkers selected by SpAn as indicated in blue in Fig. 4. Biomarker expression is an intrinsic property that, unlike correlations, does not describe relationships between different biomarkers, and therefore, is naturally expressed by an isolated node without any connecting edge.

The epithelial-stromal domain network is comprised of three dominant sub-networks associated with tumor-invading T lymphocytes,^46^ disruption in DNA mismatch repair cellular process, and the role of cancer associated fibroblasts (CAFs) in the desmoplastic microenvironment as indicated by the strong edge weight between smooth muscle actin (SMA) and collagen IV. CAFs are well known to promote EMT^47^ and the differential expression of beta-catenin and phosphorylated-MET in Fig. 6f is also consistent with the epithelial-stromal domain supporting the mesenchymal phenotype.^48^ These features have also been identified with those distinguishing consensus molecular subtype (CMS) 4 that is associated with a poor prognosis in comparison to the other 3 subtypes in the transcriptome-based classification.^12, 13^ Interestingly, the epithelial-stromal spatial domain also reveals the presence of DNA mismatch repair network that has been associated with regulation of T lymphocyte infiltration, a prominent feature of CMS1. Thus, the epithelial-stromal spatial domain associated with recurrence combines two features, where in contrast, each alone is associated with two different CMS subtypes. This theme extends to the epithelial spatial domain in Fig. 6d, where metabolic deregulation, a prominent feature of CMS3, and DNA mismatched repair, a hallmark of CMS1 are evident. The association of these two subnetworks in the epithelial domain has the potential to promote tumor cell growth while escaping immune surveillance. Finally, we observe a prominent tumor associated macrophage (TAM) network in the stromal spatial domain (Figure 6e). TAM polarization towards the M2 phenotype regulated by AKT/PTEN has been associated with poor prognosis in CRC that could result from their immunosuppressive and matrix remodeling phenotypes.^30^

### Spatial domain networks reveal domain-specific network biology of CRC-relevant pathways

We used STRING^49^ and KEGG^50^ databases to identify pathways enriched by biomarkers within each of the spatial domain networks and further corroborate their connections to prominent features in the CMS subtype classification. Figure 7 shows the pathways enriched in each of the three spatial domains, and further identifies those that are common to a majority of at least two of the three spatial domains. Since their identification is based on the spatial domain networks we computationally identified as significant for CRC recurrence prognosis, these pathways play a differentially important role in prognosis of CRC recurrence.

**Figure 7.**
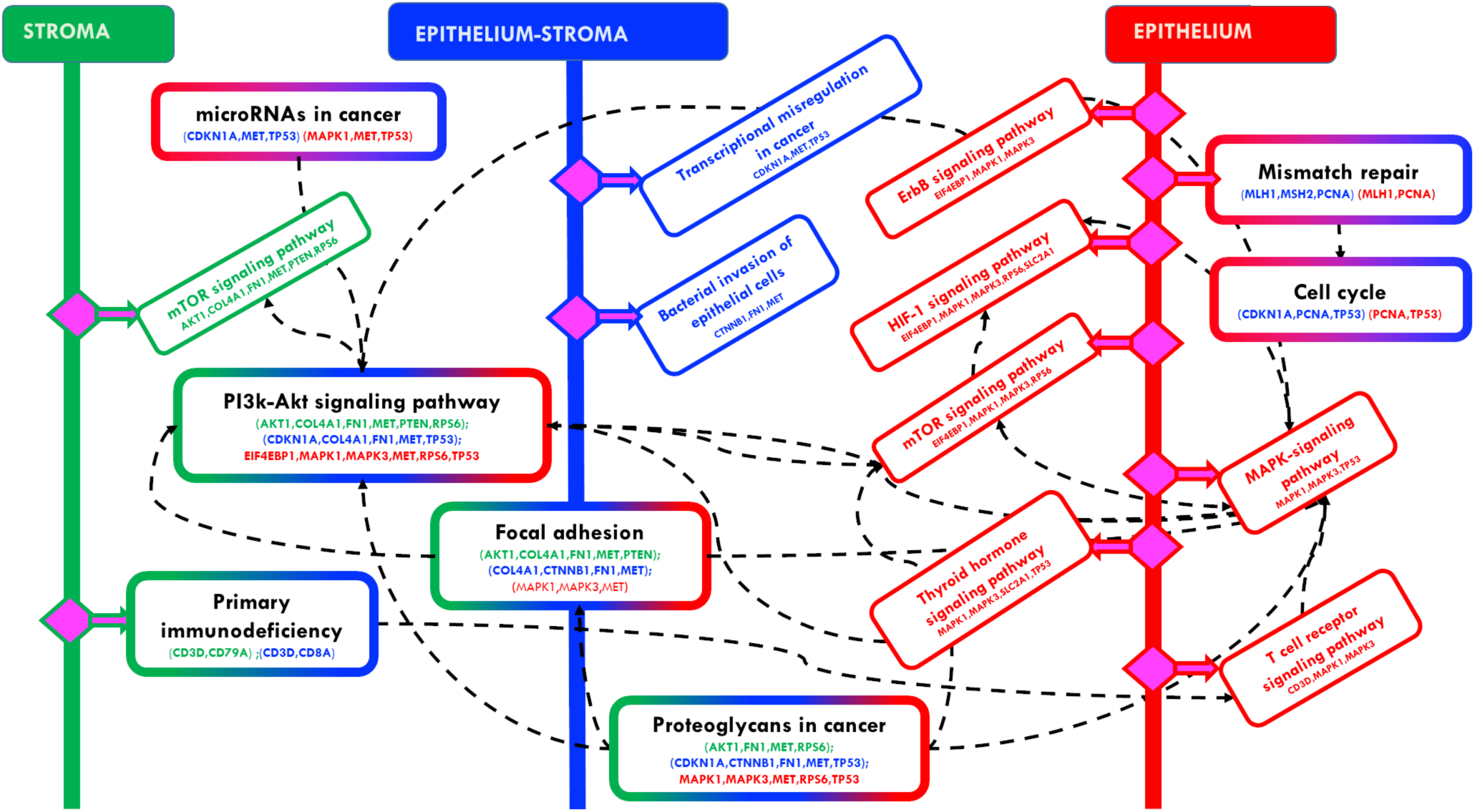
CRC recurrence-specific network biology inferred by SpAn. Domain-specific biomarkers identified by the spatial domain networks are used to interrogate the KEGG and STRING databases to identify domain-specific pathways enriched by the biomarkers. The epithelial, stromal and epithelial-stromal domain are respectively shown in green, red and blue, with pathways unique to those domains also coded with the same colors. Pathways that are enriched in more than one domain are coded with a color combination of those respective domains. For example, the PI3K-AKT signaling pathway is enriched in all three spatial-domains, and therefore, has a boundary box color-coded with all three colors. On the other hand, the mismatch repair pathway is enriched in the epithelial and epithelial-stromal domains, and is therefore, color-coded by red and blue colors.

CMS2 tumors are associated with chromosomal instability pathway and enrichment of genes associated with cell cycle and proliferation. Interestingly, both Pi3k-Akt signaling and cell cycle pathways enriched in our analysis are associated with CMS2 tumor subtype, with almost 60%-70% of CRCs associated with dysregulation of Pi3k-Akt signaling pathways.^51^

Tumors associated with the CMS4 mesenchymal phenotype show upregulated expression of genes involved in epithelial-to-mesenchymal transition along with increased stromal invasion, angiogenesis and transforming growth factor-β (TGF-β) activation.^12, 13^ Interestingly, proteoglycans in cancer, focal adhesion and microRNAs in cancer pathways enriched in our analysis enable the mesenchymal phenotype. For example, non-coding microRNAs both regulate and are targets of upstream regulators for modulating the epithelial to mesenchymal phenotype by targeting EMT-transcription factors such as ZEB1, ZEB2, or SNAIL.^52^ Similarly, the focal adhesion pathway through the integrin family of transmembrane receptors mediates attachment to the extracellular matrix, and when dysregulated promotes cell motility and the mesenchymal phenotype.^53, 54^ Furthermore, extracellular and cell surface proteoglycans with their interaction with cell surface proteins such as CD44 have been known to promote tumor cell growth and migration.^55, 56^

Our analysis suggests that by capturing correlation-based crosstalk between heterocellular signaling pathways, SpAn leverages the interconnections between the subtypes for a high performing CRC recurrence prognosis and reveals a synergistic role of the CMS subtypes in CRC progression and recurrence. We note that the ability of SpAn to leverage these interconnections is due to the spatial-context-preserving sampling of a diverse set of CRC-relevant biomarkers enabled by HxIF imaging.

Interestingly, this network biology paradigm also shows enrichment of pathways specific to a single spatial domain whose oncogenic or tumor suppressive roles in CRC is an active area of research but whose differential role in CRC recurrence has not been widely studied. For example, in the epithelial domain our analysis shows the enrichment of Thyroid hormone signaling pathway that has been associated with a tumor suppressive role in CRC development.^57, 58^ In contrast, the bacterial invasion pathway, enriched in epithelial-stromal boundary region, has been implicated in the oncogenic role of the colonic microbiome in CRC development.^59, 60^

Our analysis also reveals enrichment of certain other pathways, such as the hypoxia-inducible factor 1 (HIF-1), human epidermal growth factor receptor 2 (HER2) and T-cell receptor signaling pathways in the epithelial domain. Hypoxia is typical in many solid tumors in CRC with HIF-1 regulating tumor adaptation to hypoxic stress.^61^ Alterations in Her-2 signaling, either through genomic amplification or mutations is tumor promoting, and anti-HER2 therapies for preventing CRC recurrence and are a focus of on-going work.^62^ We finally note that MAPK and PI3K-AKT signaling cascades are implicated in many of the above discussed pathways.

## Discussion

This study highlights the importance of spatial context of the primary tumor microenvironment in conferring distinct malignant phenotypes such as recurrence in CRC. We show how a computationally unbiased approach can be implemented through statistical modeling of spatially defined domains leading to a highly specific and sensitive platform for prognostic and diagnostic tests, as well as potentially inferring therapeutic strategies (Figure 8). SpAn provides testable hypotheses regarding how the spatial association of common networks could potentially lead to emergent signaling networks conferring malignant phenotypes. For example, an epithelial domain network coupling cell metabolism and DNA repair is consistent with tumor cell growth at the expense of T cell exclusion and functional deficiency^12, 13, 63^ (Figs. 6 and 7). Likewise the hijacking of CAFs to support EMT in the context of diminished immune surveillance in the epithelial-stromal spatial domain^47, 64, 65^ (Figs. 6 and 7) and the PI3K/AKT mediated polarization of TAMs within the stromal domain (Figs. 6 and 7) can conspire to facilitate migration of tumor initiating cells to promote both local and distant recurrence.^66^

**Figure 8.**
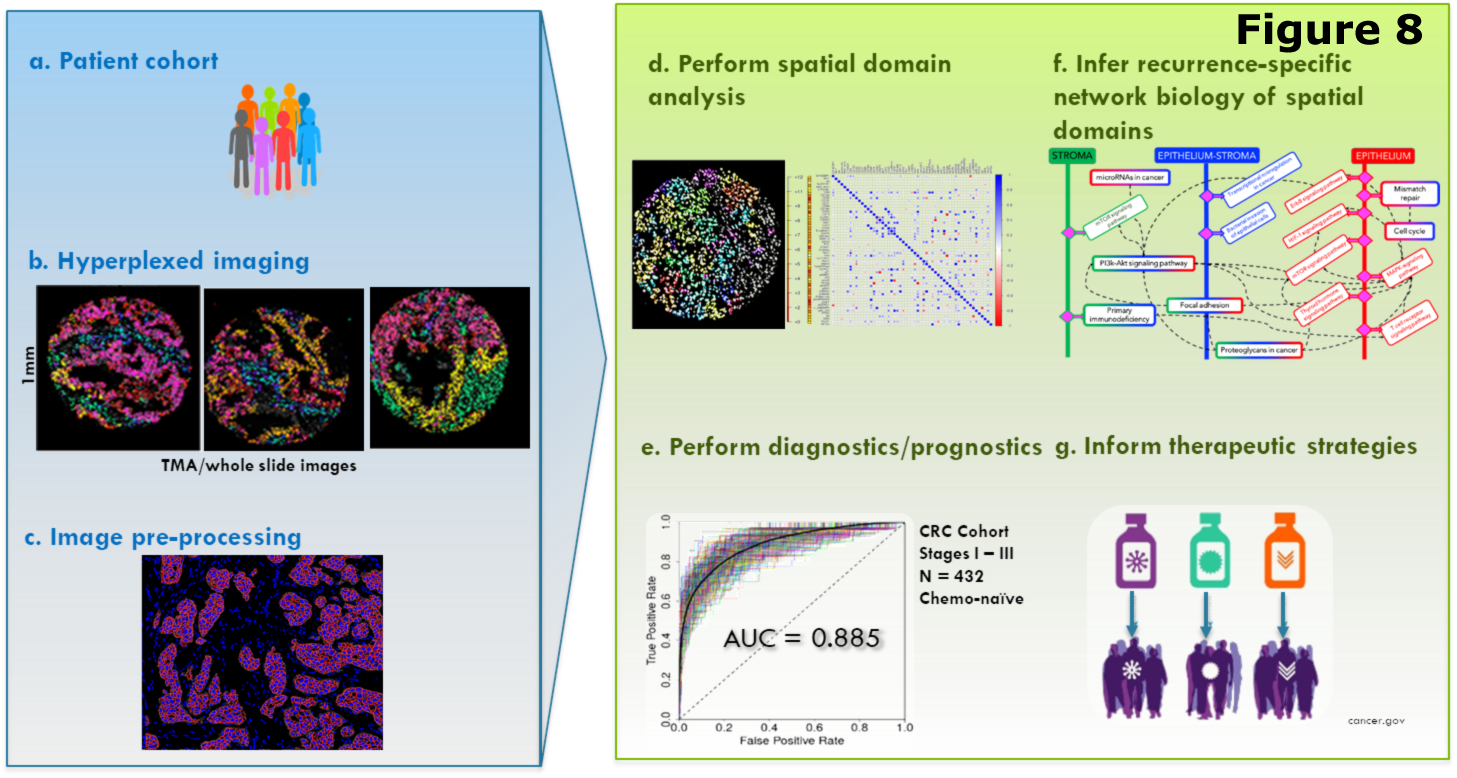
Workflow of <u>sp</u>atial <u>an</u>alytics (SpAn) computational and systems pathology platform. (a-c) SpAn utilizes images of resected primary tumors from TMAs or whole slide images based on hyperplexed fluorescence and other imaging modality platforms,^18–21^ to (d) perform spatial domain analysis for (e) patient diagnoses and prognoses and (f) infer recurrence-specific spatial domain networks to (g) potentially inform therapeutic strategies.

SpAn, when used in combination with a non-destructive hyperplexed imaging platform such as Cell DIVE^TM^ allows mechanistic hypotheses to be tested through iterative probing of the same spatial domains with additional biomarkers inferred by the pathway analyses (Figures 7 and 8). We expect even more specific mechanistic biomarkers to emerge based on the iterative hyperplexed imaging approach that incorporates finer stage-based focus, thereby reducing the total number of biomarkers needed for optimal analyses. We will be pursuing this in subsequent studies. This feature of SpAn combined with a hyperplexing imaging platform will not only allow refinement of its prognostic ability, but since the iterative analysis can be potentially conducted in real time with further advances in the technologies, it may allow the prognosis to be specific to individual patients. Importantly, by enabling the testing of mechanistic hypotheses in patient samples directly connected to a specific clinical outcome, SpAn may inform therapeutic strategies to prevent the outcome. For example, immunotherapy has shown benefits in microsatellite instable (MSI) CRC patients but remains refractory in microsatellite stable (MSS) CRC patients.^67^ By combining non-destructive hyperplexed imaging with iterative analysis, SpAn has the potential to identify spatial domain networks that are present in both MSI and MSS CRC patient cohorts and play a differential role in CRC recurrence in an MSI status-independent manner. Identification of such commonalties between MSI and MSS CRC spatial systems biology has the potential to help generate testable hypotheses for identifying therapeutic biomarkers important in biomarker-guided treatment of MSS CRC patients.^67^ [The next phase of this work will take advantage of sampling multiple regions of primary tumors with larger TMAs and/or whole side sections, exploring other spatial analytics ranging from simple to sophisticated spatial heterogeneity metrics^68^ and incorporating a combination of protein and nuclei acid biomarkers.^69^]

The ability of SpAn to exploit spatial context of the tumor makes it suitable to study cancers that progress via spatially mediated signaling interactions with their TME. SpAn, therefore, is applicable to solid tumors including sarcomas, carcinomas and lymphomas, which co-evolve with the TME^70^ of their abnormal tissue mass. We will pursue applications of SpAn to solid tumors beyond colorectal adenocarcinoma in subsequent studies. Our present retrospective study provides the foundation for such studies in other solid tumors. It also establishes feasibility of implementing SpAn in prospective studies predicting disease outcomes in patients with CRC and other malignant solid tumors. The high specificity and sensitivity of SpAn lies in its ability to unbiasedly identify emergent networks that appear to be closely associated and likely to be mechanistically linked to recurrence. We anticipate that hyperplexed datasets based on multiple imaging modalities will be generated faster and become less expensive as the technology evolves to become a mainstay tool to analyze solid tumors.

## Acknowledgements

The work of S.U. was supported in part by NIH/NCI R01CA232593. The work of D.M.S., A.G., D.L.T. and S.C.C. was supported by grant NIH/NCI U01CA204836. S.C.C. was also supported in part by NIH/NHGRI U54HG008540, and UPMC Center for Commercial Applications of Healthcare Data 711077. D.L.T. was supported in part by NIH P30CA047904, and PA DHS 4100054875. D.M.S. was supported in part by NIH/NIBIB 5T32EB009403-07. F.G was supported by NIH/NCI RO1CA208179.

## Author Contributions

S.U., A.M.S., D.L.T. and S.C.C. conceived and designed the study and wrote the manuscript. S. U. and S.C.C. developed the computational platform. S.U. performed the formal analyses. S.U., A.M.S., F.P., D.L.T. and S.C.C developed the systems biology framework. C.J.S and F.G. selected and validated the biomarkers, performed imaging and acquired the data, quality checks and preliminary data analysis. S.U., C.J.S., S.F., F.P., D.M.S. and L.N. were responsible for data quality control and providing analysis support. A.G., C.J.S. and F.G. helped with technical issues and edited and reviewed the manuscript.

## Acknowledgments

The authors are grateful to the following team members for their early role in development of this work: Yunxia Sui and Brian Ring for assembling and analyzing the multiplexed data, Alberto Santamaria-Pang and Yousef al-Kofahi for single cell segmentation; Alex Corwin for supporting Cell DIVE acquisition software.

## Methods

### Patient Cohort and Tissue microarray (TMA)

As detailed in Gerdes et al.,^1^ the CRC cohort in this analysis was collected from the Clearview Cancer Institute of Huntsville Alabama from 1993 until 2002 with 747 patient tumor samples collected as paraffin embedded specimens. Tissue microarrays were constructed by Applied Genomics (later part of Clarient Lab). to facilitate large scale biomarker analysis. Cores with 0.6 mm diameters from the patient samples were distributed across seven slides. After quality control measures were taken, 694 TMA patient spots remained for analysis. Sample attrition was due to insufficient tumor fraction in the representative TMA core. Of the remaining samples, 450 were chemo-naïve CRC patients that were treated with surgery alone, and the remaining 244 patients were treated with 5-fluorouracil based chemotherapy regimens. 432 chemo-naïve patients were used in this study. Supplementary Table S2 details the median age, gender, recurrence, recurrence time and survival times for the 432 Stage I-III CRC patients. As can be seen the patient cohort is balanced in age and gender across the three stages.

### Antibody validation

We ensured that the correct biomarker expression was captured by the imaging system using an antibody standardization process.^1^ Specifically, antibodies were selected based on their staining specificity and sensitivity, compatibility with the two-step antigen retrieval, and resilience during 1, 5, and 10 rounds of dye inactivation chemistry. Depending on the marker, a variety of specificity tests were conducted including, immunogen peptide blocking before incubation with tissue, drug-treated fixed cell lines, fixed cell lines with gene amplification or deletion, phosphatase treatment of samples to verify phospho-specificity, and visual inspection by expert pathologists of expected localization patterns. Furthermore, fluorescent dyes were conjugated to the primary antibody at several initial dye substitution ratios and specificity of each conjugate was verified and sensitivity compared with levels found in previous experiments. Staining performance was assessed by expert biologists and poor or non-specific staining was excluded.

### Cell DIVE-based hyperplexed imaging (HxIF) of tissue microarrays (TMA)

The 55 biomarkers plus DAPI nuclear counterstain included in this study are described in Fig. 1 and Supplementary Table S1. HxIF imaging of a TMA slide was performed using sequentially multiplexed labeling and imaging of 2 to 3 biomarkers along with DAPI counterstain through a label–image–chemical-inactivation iterative cycle visualized in Supplementary Fig. S1, and previously described^1^ in detail in the supporting information therein. Broadly, the supporting information details the hyperplexed immunofluorescent workflow with information on iterative cycles of antibody labeling of single 5 µm formalin-fixed and paraffin embedded tissue sections and TMA slides, autofluorescence removal, imaging, and dye inactivation in tissue. All samples were stained and imaged in a single batch for 2 to 3 biomarkers and DAPI at a time.

### Image processing and single cell analysis

DAPI based nuclear staining was used to register and align sequentially labeled and imaged TMA spots prior to downstream image analysis steps.^1, 2^ Autofluorescence was removed from the stained images,^1, 3, 4^ which were then segmented into epithelial and stromal regions (Fig. S2), differentiated by epithelial E-cadherin staining. This was followed by segmentation of individual cells in both the epithelium and stroma. Epithelial cells were segmented using Na^+^K^+^ATPase based cell-membrane staining to delineate cell borders and membrane regions, the cytoplasmic ribosomal protein S6 for cytoplasm identification, and DAPI stain for nuclear regions.Protein expression level and standard deviation were subsequently quantified in each cell.

### Quality checks and data normalization

Following single cell segmentation, several data pre-processing steps were conducted. These included cell filtering, spot exclusion, log2 transformation and slide to slide normalization. Cells were included for downstream analysis if their size was greater than 10 pixels at 20X magnification. The hyperplexing process can result in the tissue being damaged, folded or lost. Image registration issues can also result in poor-quality cell data. Therefore, a tissue quality index based on the correlation of that image with DAPI was calculated for each cell for each round. Only those cells whose quality index equals to or greater than 0.9 (meaning that at least 90% of the cells overlapped with DAPI) were included. All the slides for all the biomarkers were adjusted to a common exposure time per channel. The data were then log2 transformed. A median normalization that equalizes the medians of all the slides was performed to remove slide to slide non-biological variability.

### SpAn Input Features

For each of the epithelial, stromal and epithelial-stromal spatial domains, SpAn used *M* = 1540 domain-specific biomarker feature vector ***f*** as input. This input feature vector comprised of (1) mean intensity value of 55 biomarkers averaged across all cells within the spatial domain, and (2) 1485 (= 55*54/2) Kendall rank-correlations between all 55 biomarker pairs. Kendall rank-correlation was chosen as the correlation metric because it is a non-parametric measure of association between two biomarkers. Moreover, its use of concordant and discordant pairs of rank-ordered biomarker expression for computing correlation coefficients allows it to robustly capture biomarker associations in presence of measurement noise and small sample size. Rank-correlation for each pair of biomarkers was computed for each spatial-domain from all cells across the spatial-domain expressing the biomarkers. This approach is distinctly different from prediction models that typically consider correlations via interactions, implicit within the models, between mean biomarker intensity expressions – with the biomarker expressions being the only covariates of the model.^5^ We emphasize that we did *not* compute correlations through mean intensity biomarker expression across the spatial-domain, but instead used biomarker expressions across individual cells of the spatial domain to explicitly compute domain-specific rank-correlation values between every pair of biomarkers to form the SpAn correlation feature set.

Before computing these two sets of features, SpAn analysis workflow included an initial intensity threshold step to ensure feature robustness. Specifically, we computed intensity-based distribution of cell-level biomarker expression separately for every biomarker across each patient TMA spot. Only intensities above the 85^th^ percentile on this distribution were considered as biomarker expression and included in computing the intensity features. This focus on the right-tail of the intensity distribution was deliberately conservative, and although it might have potentially excluded low-intensity biomarker expression, it minimized inclusion of false-positive expressions into the analysis.

### Penalized Cox proportional hazard regression

For each spatial-domain, SpAn implemented the Cox proportional hazard model via the partial likelihood function 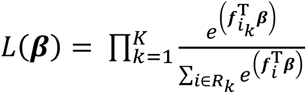 with the penalty given by 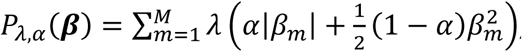, and α = {0,1}. (The validity of using the Cox proportional hazard regression model is demonstrated in Fig. S3.) Given feature vector ***f*** as input, the partial likelihood *L*(*β*) quantifies the conditional probability of observing CRC recur in a patient at time *t*_*k*_(proportional to the numerator 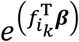 of *L*(*β*)), given the risk that a patient will recur from the set *R*_*k*_ of patients at risk at time *t*_*k*_(proportional to the denominator 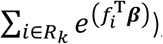, over all time *t*_*k*_, *k* = 1, …, *K*, as quantified by the product over time index *k*. The partial likelihood is a function of the coefficient vector *β*, whose penalized estimate is then used to compute the proportional hazard ratio *HR* = *e*^(***f***^T^*β*)^. SpAn computed this estimate via a two-step process that first selects the parsimonious set of features required for optimally predicting the risk of recurrence, and then learns the model predicting the risk of recurrence based on the selected features.

The feature selection step is implemented via L1-penalized (LASSO) Cox regression where *α* is set to 1 in penalty *P*_*λ*,*α*_(*β*). LASSO based L1-penalized model selection performs feature selection by forcing the coefficients of vector *β* that play a minimal role in predicting risk of recurrence to zero. This is done in a principled manner by minimizing the model deviance along the LASSO regularization path.^6, 7^ The features corresponding to the non-zero coefficients in *β* are the features selected by SpAn to define the final functional form of Cox proportional hazard model. Model learning based on this functional form is performed in the second step via maximizing the partial likelihood function with L2-regularization as the penalty, implemented by setting *α* to 0 in the penalty term.^6, 7^ L2-regularization allows SpAn to learn the Cox proportional hazard model while avoiding over-fitting. An advantage of this two-step process is the decoupling of feature selection from estimation of beta coefficient values, resulting in the latter not being conditioned on the complete set of 1540 features but being dependent only on the selected features.

To ensure the stability of the selected features, SpAn repeated model selection over 500 bootstraps, and included only those features that were consistently concordant at the 90% level with the recurrence outcome. (The rationale for 90% concordance is discussed in supplementary Fig. S4.) SpAn next performed a stability check on the beta-coefficients estimated in the second step. Specifically, the stability of the coefficient sign in 90% of the 500 bootstrap runs was tested, and only features that passed this threshold (Fig. 2c) were included in the spatial domain model. SpAn performed this process independently for each of the three spatial-domains resulting in domain-specific recurrence-guided features (Fig. 2c) and their coefficients (Fig. S5).

### SpAn is computationally unbiased

SpAn begins penalized Cox proportional hazard regression by including the full 1540 features. It then utilizes LASSO based shrinkage to parsimoniously optimize the full model along the L1 regularization path by minimizing model deviance.^7^ By combining this principled shrinkage via L1-penalized Cox proportional hazard regression, with bootstrapping to establish the stability of the selected subset of features at 90% concordance with the recurrence outcome (supplementary Fig. S4), SpAn avoids typical biases associated with many model selection approaches based on stepwise variable selection, backward elimination and forward selection.^8^ These biases include *R*^2^ values being biased high, *F* and *χ*^2^test-statistics not having their associated distributions, p-values being biased towards zero, and standard errors of regression coefficient estimates being biased low, while absolute values of regression coefficients being biased high.

### Spatial-Model

Each of the three recurrence-guided domain-specific models defined a hazard risk given by 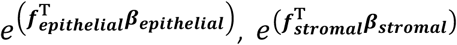 and 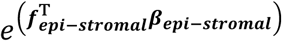 for the epithelial, stromal, and epithelial-stromal domains respectively. SpAn then defined the final overall risk of recurrence model as 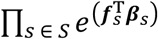, with *S* = {*epithelial, stromal, epi − stromal*}.

### Partial correlations and spatial-domain networks

For each spatial domain, the selected features identified a set of biomarkers specific to predicting risk of CRC recurrence. SpAn used them to define a space of biomarkers within which *partial* correlations between every pair was computed by controlling for confounding effect of biomarkers not defining the pair.^9^ The process performed on each patient was as follows: Let the set of biomarkers identified by the selected features be N (<= 55). Using the already computed Kendall rank-correlations between the 55 biomarkers, an *N* × *N* correlation matrix *C* corresponding to the *N* biomarkers was constructed, with small shrinkage-based modification to guarantee its positive definiteness, and therefore, its invertibility. Next, the *N* × *N* precision matrix *P* was computed by inverting *C*. The partial correlation between any two biomarkers *bm*_*i*_ and *bm*_*j*_ within the set identified by the selected features, was then computed using 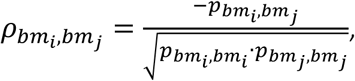, where *pbm*_*i*_,*bm*_*j*_ is the (*i*, *j*)^th^ element of the precision matrix *C*. The partial correlations were performed for all patients and were then separated into two groups corresponding to patients with no evidence of disease and those patients in which CRC recurred. Probability distributions of the partial correlations – on the compact set [−1,1] – within each group were computed and the information distance between these two distributions was computed using the Jensen-Shannon divergence. This information distance defines the differential change in the association – partial correlation – between biomarkers *bm*_*i*_ and *bm*_*j*_ in the two patient cohorts. Greater the distance, larger the differential change. Repeating this process for all *N*(*N* − 1)/2 biomarker pairs resulted in the information distance matrices shown in Figs. 4a-c for the three spatial domains. These information distance matrices were thresholded at the 99^th^ percentile resulting in the computationally inferred spatial-domain networks shown in Figs. 4d-f. The high percentile was chosen to ensure that most discriminative networks are captured.

### Enrichment analysis

The STRING database^10, 11^ was queried with the set of proteins identified by the spatial domain networks generated by thresholding the domain-specific information distance matrices at the 99^th^ percentile, to perform functional enrichment analysis using Fisher’s exact test with multiple testing correction.

---

**Table S1:**
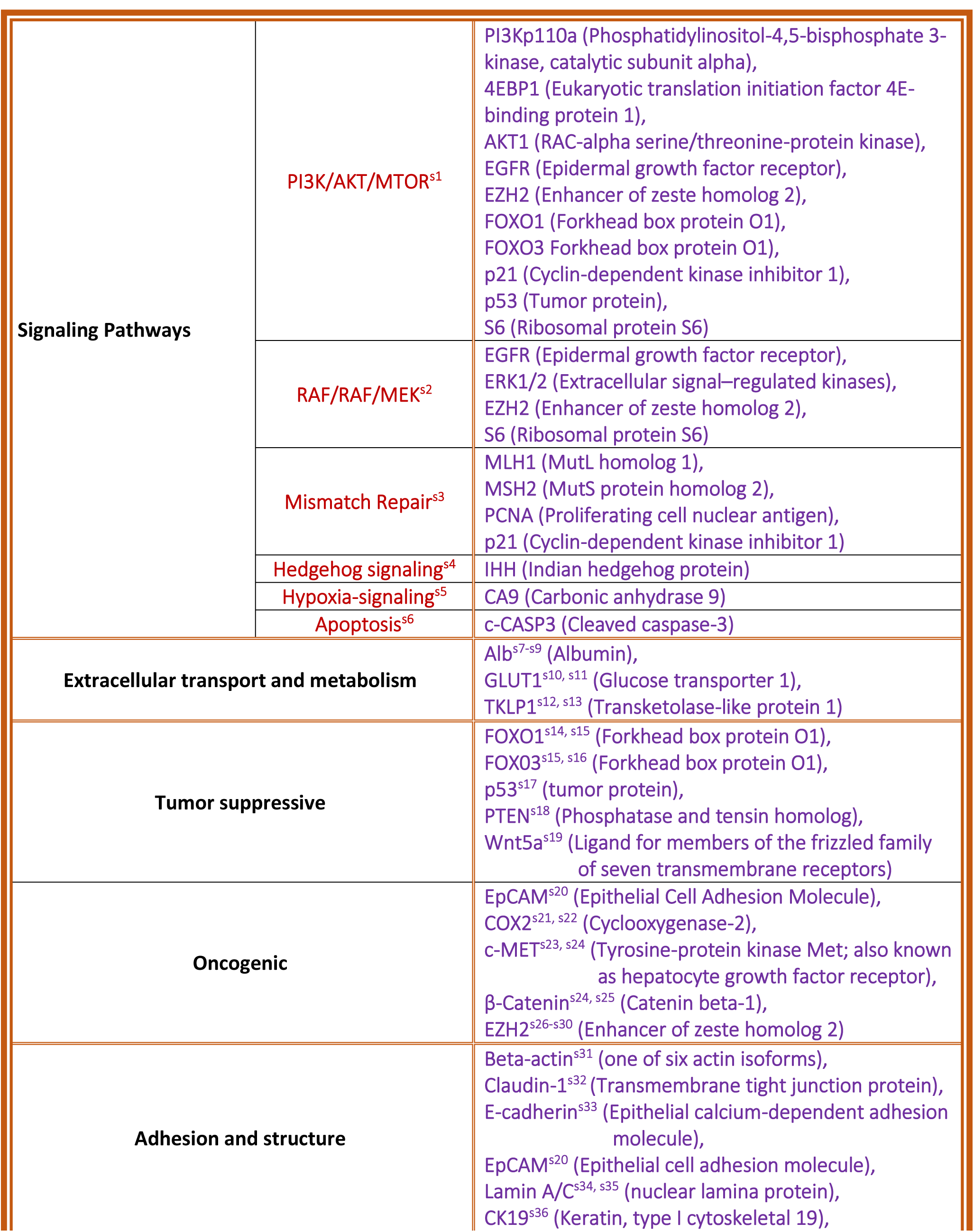
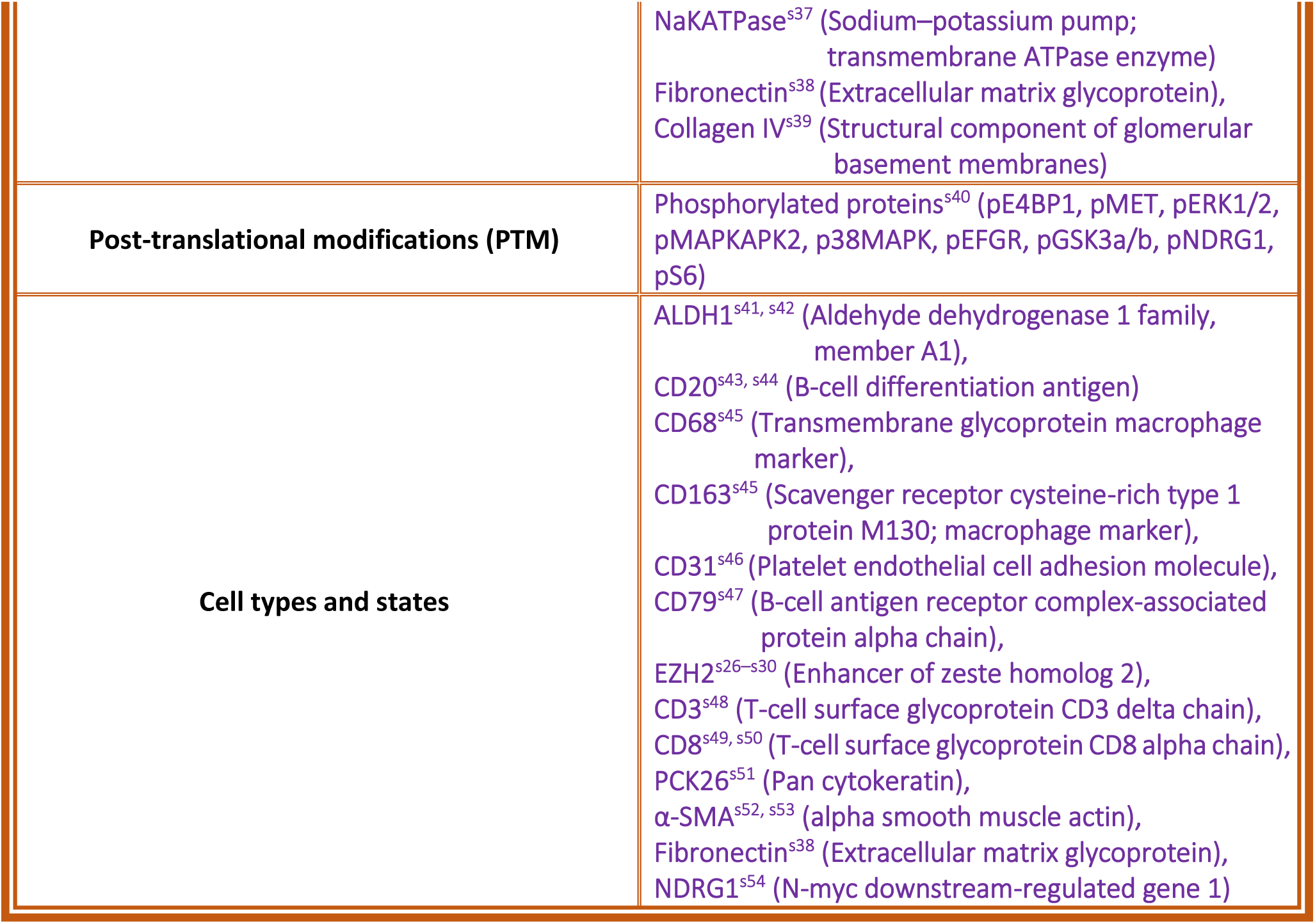
Biomarkers and their relation to colon cancer

**Table S2:**
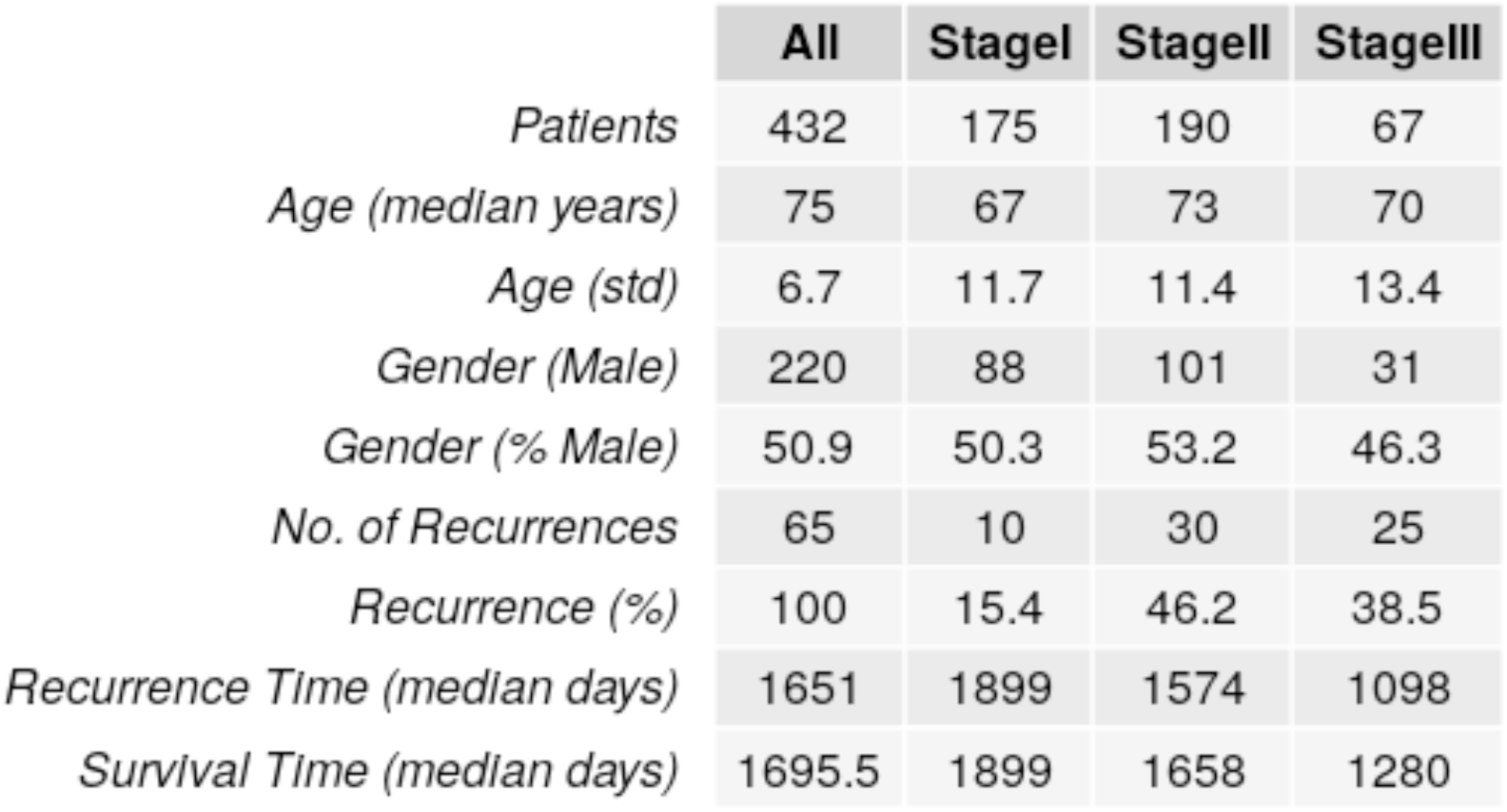
Patient Cohort and Clinical Properties

**Table S3:** List of spatial-domain features (biomarkers and their correlations) selected by SpAn

**Figure S3**
| Epithelial.spatial.domain | Stromal.spatial.domain | Epithelial.Stromal.spatial.domain |
| --- | --- | --- |
| 4EBP1.CD68 | 4EBP1.CD8 | ALDH1.CD68 |
| 4EBP1.CD79 | Akt1.pEGFR | beta-actin.NDRG1 |
| 4EBP1.pS6 | Akt1.pGSK3a | BetaCatenin.Claudin1 |
| Akt1.PCNA | beta-actin.EZH2 | BetaCatenin.pGSK3a |
| Albumin.EZH2 | CA9.CD79 | CD20.CollIV |
| ALDH1.BetaCatenin | CD163.PTEN | CD3.CD8 |
| ALDH1.NDRG1 | CD20.Fibronectin | CD3.ERK |
| CD31.CD8 | CD3.p_p38MAPK | CD68.SMA |
| CD79.p53 | CD31.CollIV | CD8.PCNA |
| CD79.pGSK3b | CD68.Fibronectin | CK19.pGSK3b |
| CD8.SMA | CD79.c-CASP3 | c-MET.NaKATPase |
| CK19.LaminA/C | CD79.pMAPKAPK2 | CollIV.SMA |
| Claudin1.Glut1 | c-MET.pNDRG1 | COX2.E-cadherin |
| E-cadherin.PCNA | CollIV.ERK | E-cadherin.NaKATPase |
| ERK.Fibronectin | CollIV.pNDRG1 | E-cadherin.PCNA |
| ERK.pGSK3b | ERK.LaminA/C | EGFR.pMAPKAPK2 |
| Fibronectin.pGSK3b | Fibronectin.PI3Kp110a | MLH1.MSH2 |
| FOXO1.IHH | Glut1.SMA | NaKATPase.pGSK3b |
| Glut1.PTEN | LaminA/C.pERK | S6.Wnt5a |
| IHH.pMET | LaminA/C.PTEN | BetaCatenin |
| MLH1.pMET | pEGFR.p_p38MAPK | c-MET |
| NaKATPase.p_p38MAPK | pS6.SMA | CollIV |
| NDRG1.p_p38MAPK | CD3 | Fibronectin |
| pERK.pMET | c-MET | IHH |
| pMAPKAPK2.pS6 | IHH | p21 |
| CD3 | pGSK3a | p53 |
|  | S6 | pGSK3a |
|  |  | pMET |

**Figure S1.**
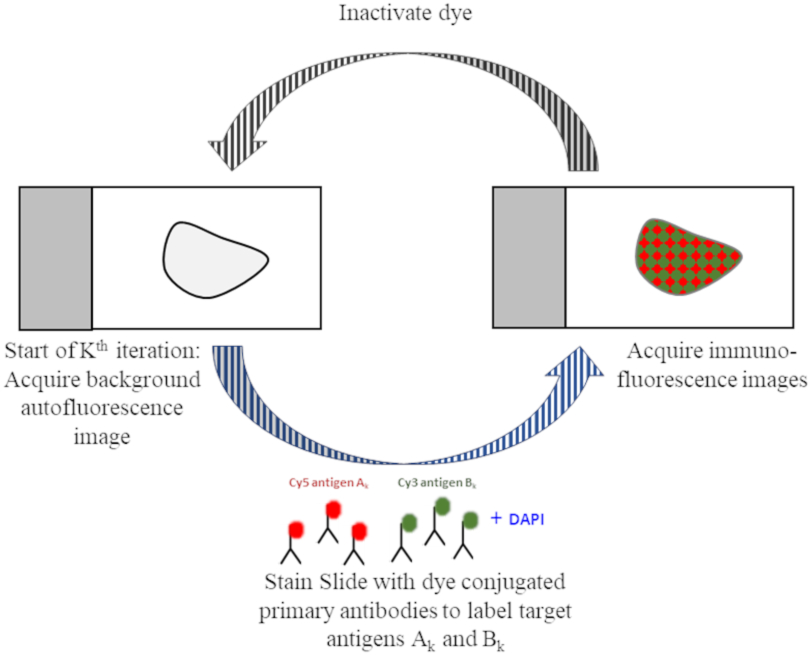
Cell DIVE hyperplexed immunofluorescence imaging and processing scheme. For the K^th^ iteration, autofluorescence image of the tissue is acquired prior to labeling it with 2-3 fluorescent dye-conjugated primary antibodies and DAPI counterstain. Fluorescent-labeled tissue images are then acquired, followed by inactivation of the dyes and start of the next iteration.^20^

**Figure S2.**
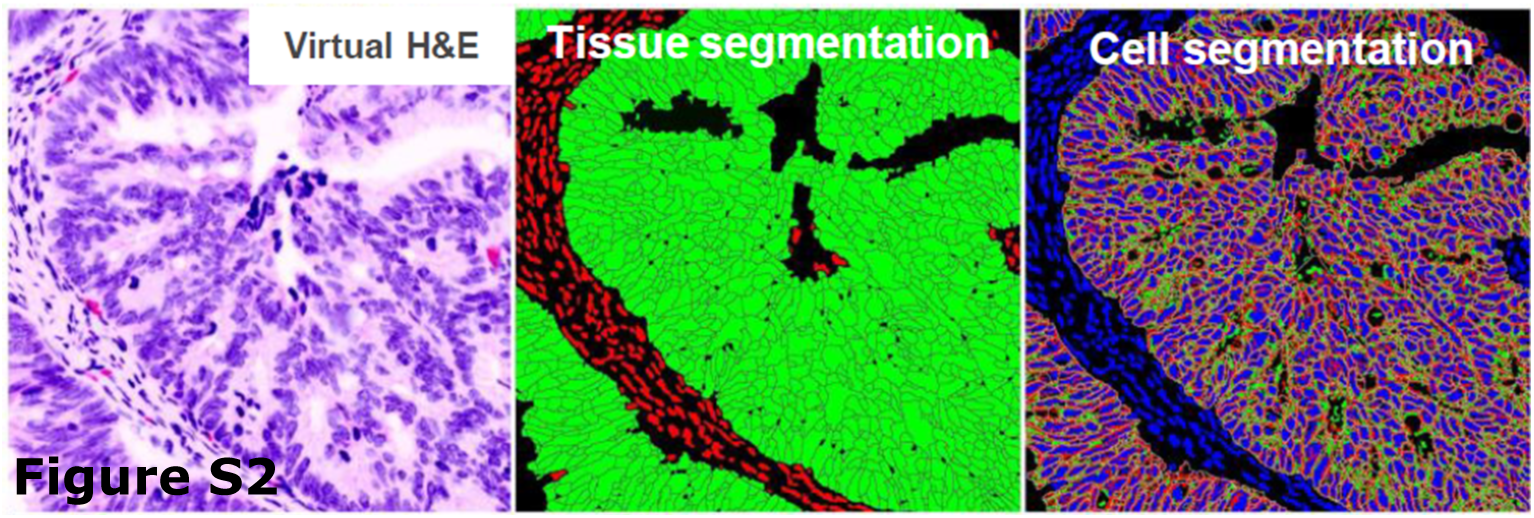
Tissue and cell segmentation. TMA spot visualized here through a virtual Hematoxylin and Eosin image. Tissue segmentation is performed by using expression of E-cadherin, a highly epithelial cell-specific marker, to identify the epithelial spatial domain, shown in green. The remaining region of the TMA spot is identified as the stromal spatial domain, shown in red. The epithelial-stromal spatial-domain runs all along the boundary between the epithelial (green) and stromal (red) spatial-domains and has a width of 100 µm. Individual cell segmentation in the epithelial spatial domain is performed using expression of Na^+^K^+^ATPase (cell membrane marker), ribosomal protein S6 (cytoplasmic marker) and DAPI (nuclear counterstain, shown in blue). The remaining cells are assigned to the stromal spatial domain.

**Figure S3.**
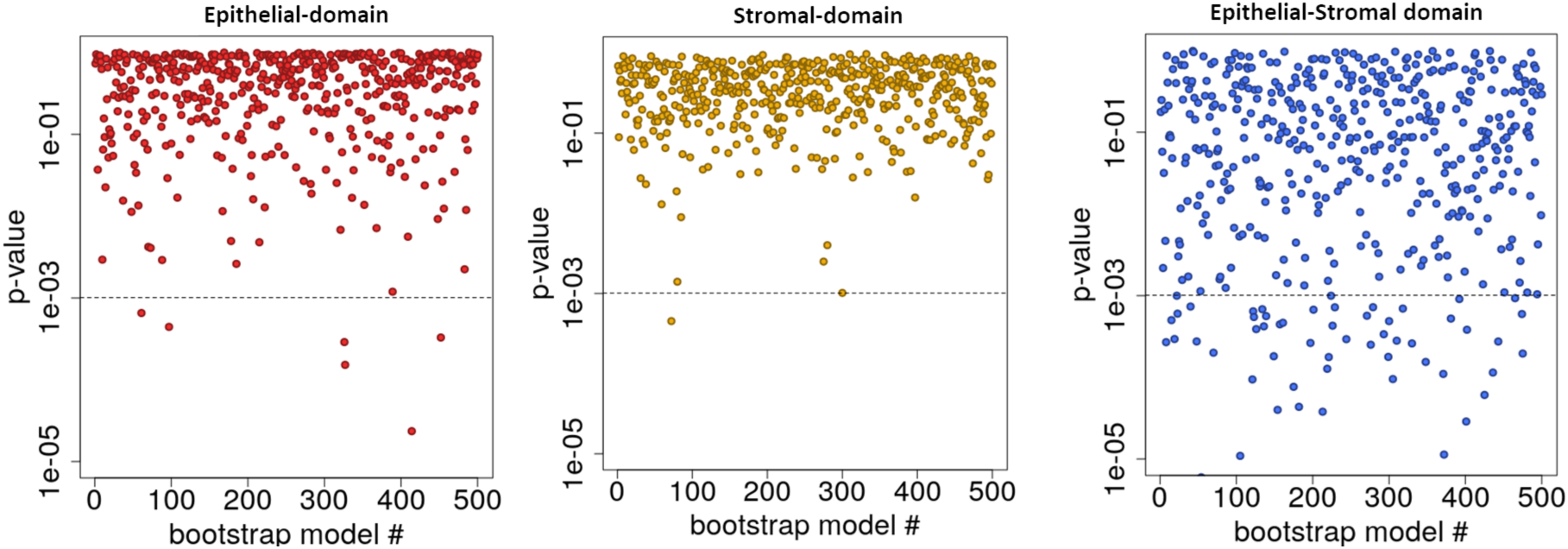
Validity of the proportional hazard assumption in penalized Cox regression. The p-values (shown in log scale) measure the significance of the relationship between scaled Schoenfeld residuals and time to recurrence. A non-significant relationship between the two indicates the validity of the proportional hazard assumption for the overall Cox regression. It can be seen that for each of the three domains, the overall global test is not statistically significant at the 95% confidence level (indicated by the dashed line) for almost all Cox models generated for the 500 bootstrap runs, demonstrating that the proportional hazard assumption is consistently valid. The p-values were computed using the cox.zph function in the *survival* R package.

**Figure S4.**
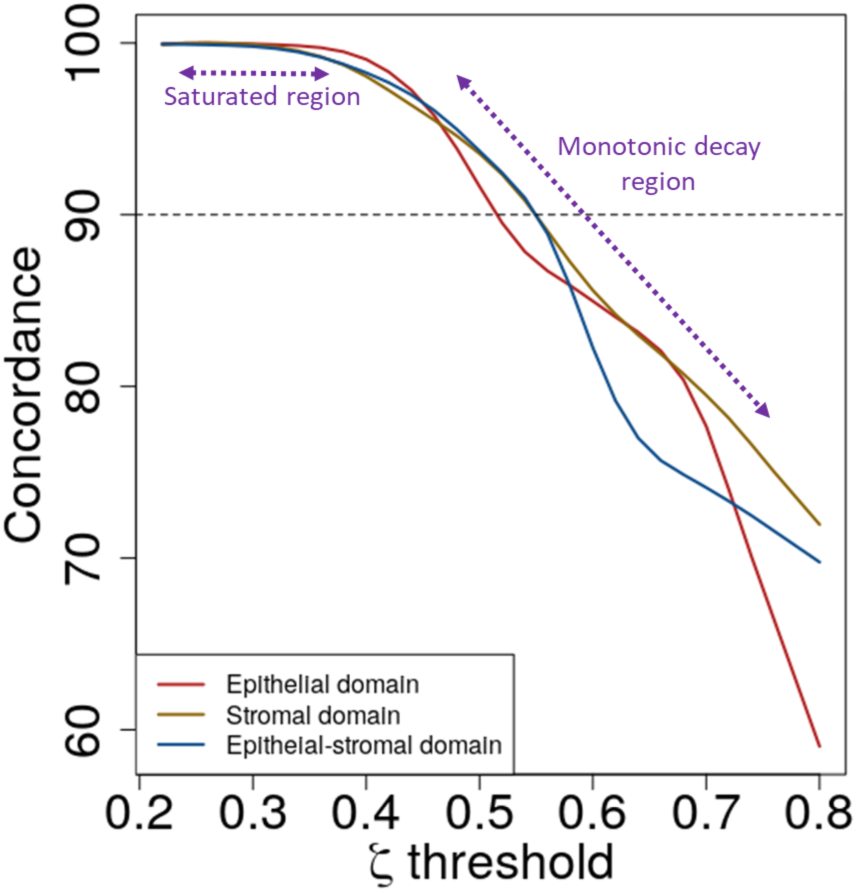
Rationale for choosing 90% concordance rate. Plot of concordance of the penalized Cox regression model as a function of a threshold function that identifies the biomarker features most consistently selected by L1 penalization at the concordance level corresponding to the threshold. The larger the threshold the more stringent the consistency requirement on feature selection, and smaller the number of selected features. As shown in the plot, for low threshold values, the concordance value is saturated, and therefore, in this region injective correspondence between threshold value and concordance does not exist. In the monotonic decay region such a correspondence can be identified. The 90% concordance level identifies such a correspondence for all three spatial domains without compromising performance.

**Figure S5.**
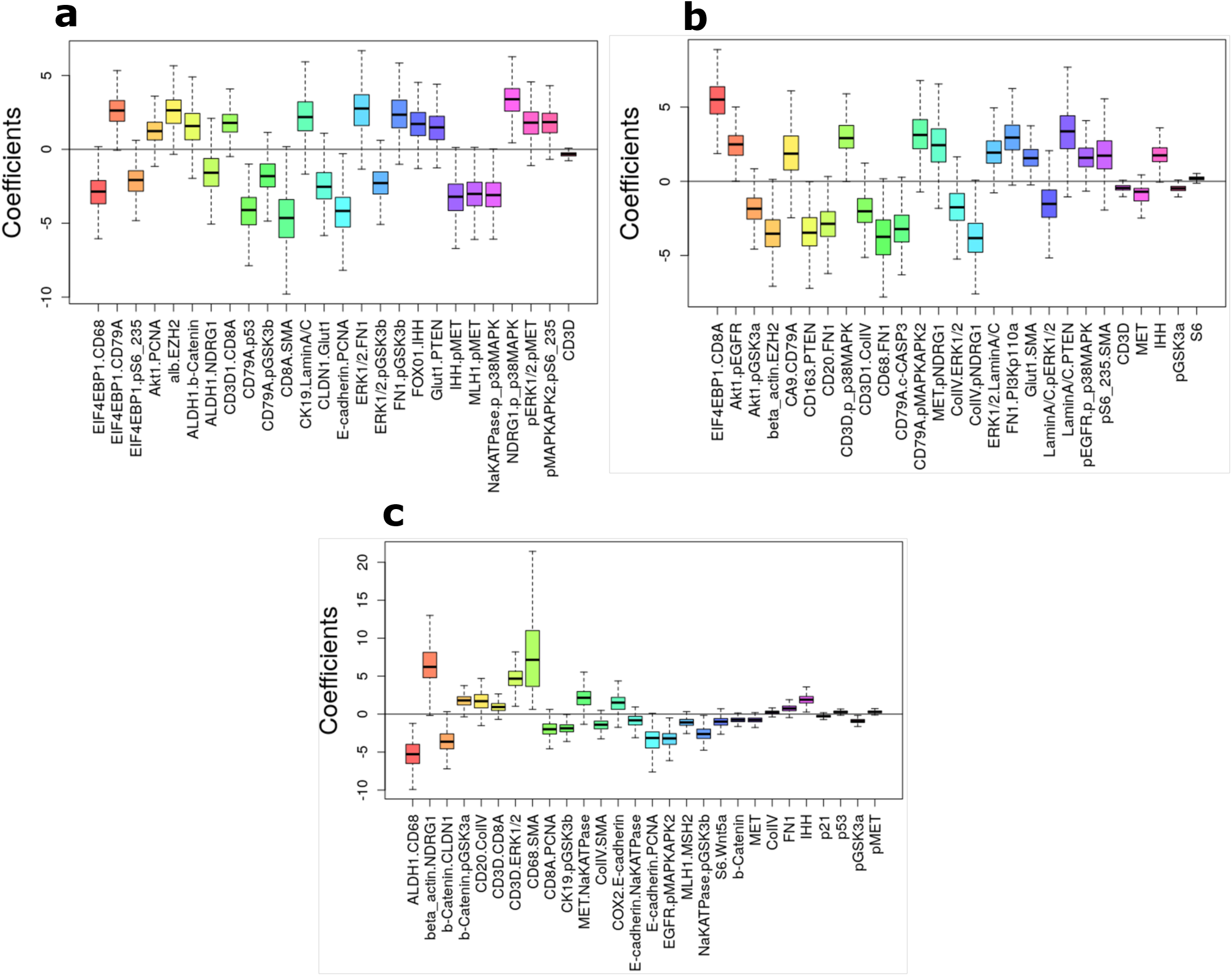
Coefficients for the recurrence-guided and domain-specific penalized Cox regression models of SpAn. Boxplots for coefficients that control the contribution of the selected features (obtained using L1-penalty) to each of the recurrence-guided and domain-specific penalized Cox regression under L2 regularization. The coefficients were computed for all 500 bootstrap runs and the boxplots capture the spread of values. It is worth noting that for all bootstraps the coefficients maintain their sign, which quantifies the nature of their contribution. A positive coefficient implies worse prognosis for increase in the corresponding feature value, while negative coefficient implies the converse.

**Figure S6.**
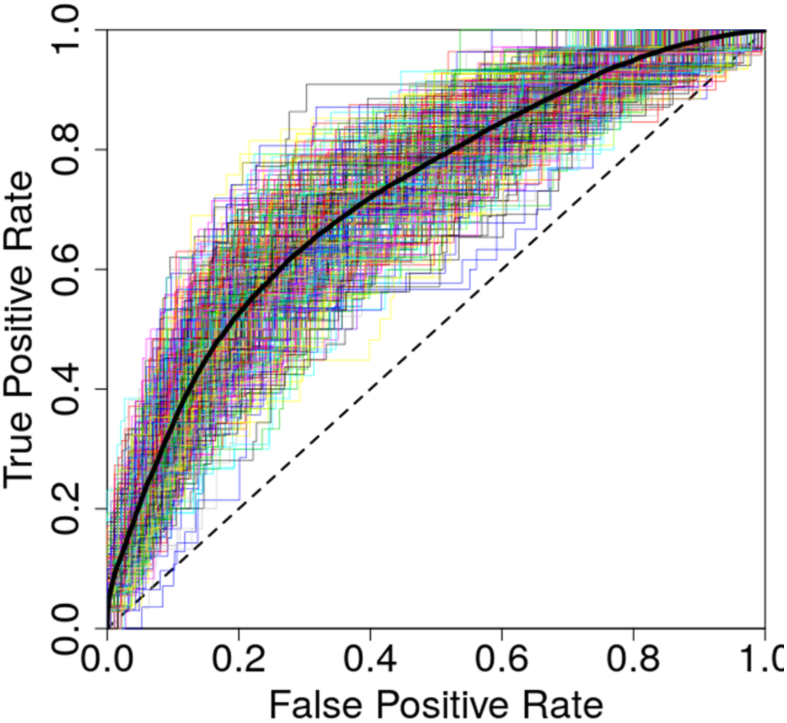
Performance of SpAn platform for predicting risk of 5-year CRC recurrence for intensity-based features. SpAn ROC curves for predicting risk of 5-year CRC recurrence in patients with resected CRC primary tumor using only biomarker expressions. The plot shows ROC curves for 500 bootstrap runs with independent training and validation sets. Mean area under the ROC curve is 72% with a standard error of 0.2%.

**Figure S7.**
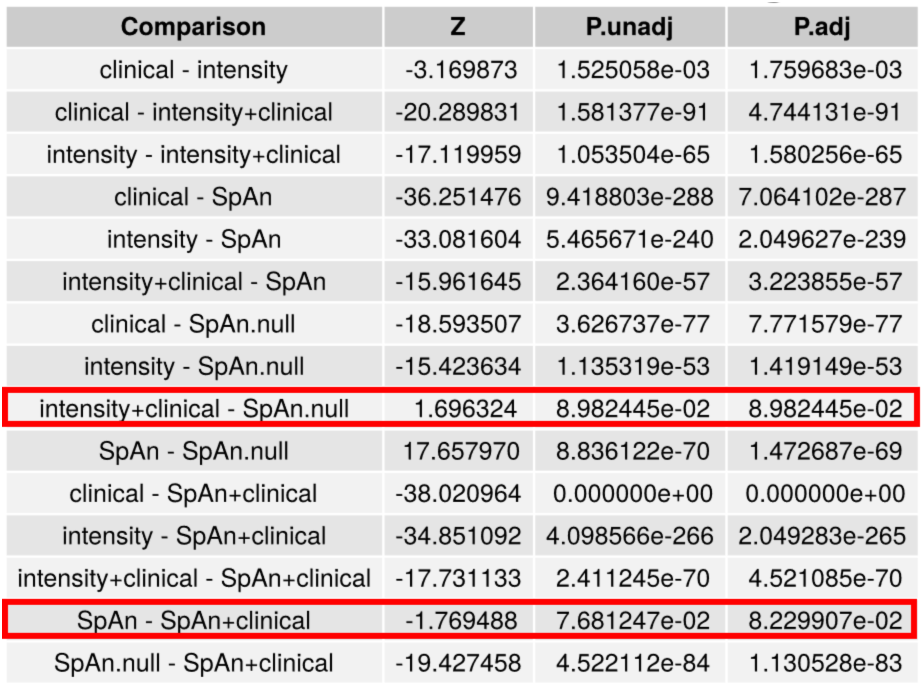
Dunn’s pairwise multiple comparison post-hoc analysis between prediction models based on non-parametric Kruskal-Wallis test. Statistical significance of pairwise performance comparison between SpAn and five other prediction models that include clinical model, biomarker expression model (denoted by intensity), SpAn.null model (denoting SpAn without spatial-domain context), biomarker expression + clinical model and SpAn + clinical model. All pairwise difference in performance are statistically significant at the 99% confidence interval expect difference between 1. biomarker expression + clinical and null models, and 2. SpAn and SpAn + clinical models.

**Figure S8.**
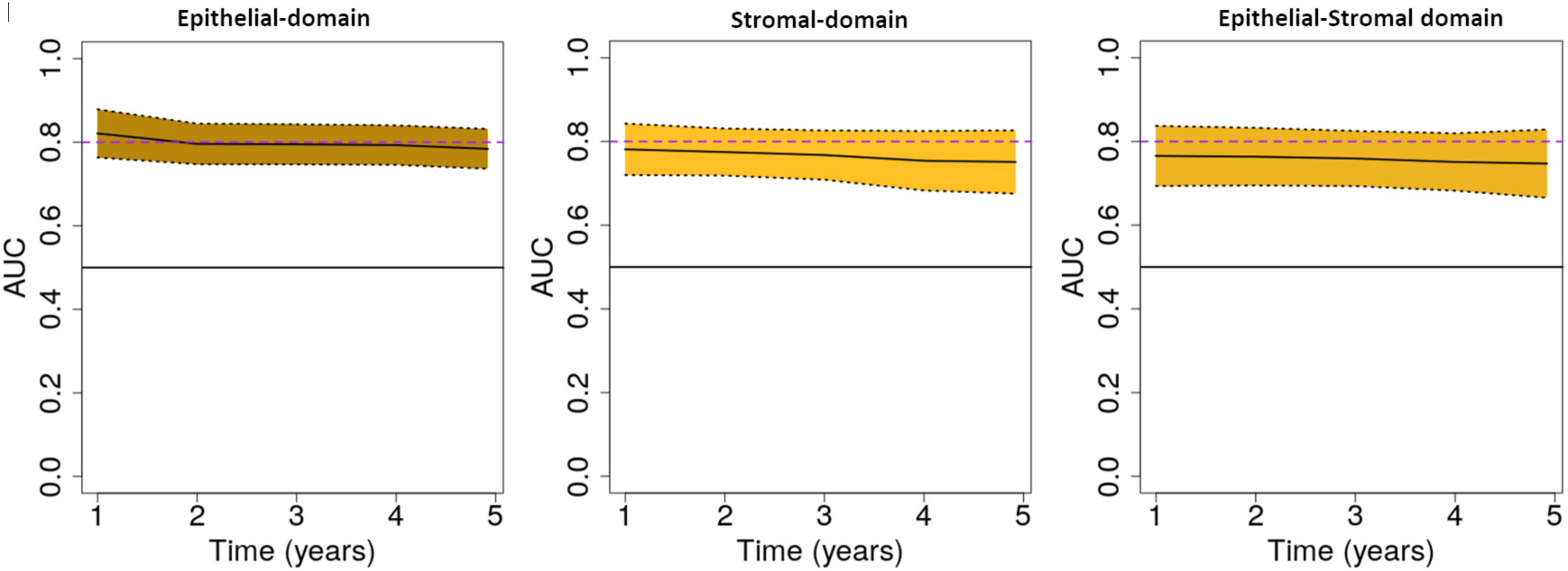
Time-dependent AUCs for domain-specific temporal performance of SpAn. Temporal performance of SpAn for the three spatial-domains illustrated by the time-dependent AUC values plotted as a function of time in years. The 95% confidence interval computed using the 500 bootstraps for each of the three spatial-domains is also shown by the yellow shaded area around the mean time-dependent AUC values depicted by the solid black line.

## References

1. Bray, F., et al. Global cancer statistics 2018: GLOBOCAN estimates of incidence and mortality worldwide for 36 cancers in 185 countries. CA. Cancer J. Clin. 68, 394–424 (2018).

2. Hanahan, D. & Weinberg, R. A. Hallmarks of Cancer: The Next Generation. Cell 144, 646– 674 (2011).

3. Weber, C. E. & Kuo, P. C. The tumor microenvironment. Surg. Oncol. 21, 172–177 (2012).

4. Tape, C. J. The Heterocellular Emergence of Colorectal Cancer. Trends in cancer 3, 79–88 (2017).

5. Weiser, M. R. AJCC 8th Edition: Colorectal Cancer. Ann. Surg. Oncol. 25, 1454–1455 (2018).

6. Gospodarowicz, M. K., Brierley, J. D. & Wittekind, C. TNM Classification of Malignant Tumours. (Wiley, 2017).

7. Mlecnik, B., Bindea, G., Pagès, F. & Galon, J. Tumor immunosurveillance in human cancers. Cancer Metastasis Rev. 30, 5–12

8. Bir, A. S., Fora, A. A., Levea, C. & Fakih, M. G. Spontaneous Regression of Colorectal Cancer Metastatic to Retroperitoneal Lymph Nodes. Anticancer Res. 29, 465–468 (2009).

9. Stanta, G. & Bonin, S. Overview on Clinical Relevance of Intra-Tumor Heterogeneity. Frontiers in Medicine 5, 85 (2018).

10. Carmona-Fontaine, C. et al. Metabolic origins of spatial organization in the tumor microenvironment. Proc. Natl. Acad. Sci. 114, 2934 LP – 2939 (2017).

11. Marusyk, A. et al. Spatial proximity to fibroblasts impacts molecular features and therapeutic sensitivity of breast cancer cells influencing clinical outcomes. Cancer Res. canres.1457.2016 (2016). doi:10.1158/0008-5472.CAN-16-1457

12. Guinney, J. et al. The consensus molecular subtypes of colorectal cancer. Nat. Med. 21, 1350–1356 (2015).

13. Dienstmann, R. et al. Consensus molecular subtypes and the evolution of precision medicine in colorectal cancer. Nat. Rev. Cancer 17, 79 (2017).

14. Tauriello, D. V. F., Calon, A., Lonardo, E. & Batlle, E. Determinants of metastatic competency in colorectal cancer. Mol. Oncol. 11, 97–119 (2017).

15. Pagès, F. et al. International validation of the consensus Immunoscore for the classification of colon cancer: a prognostic and accuracy study. Lancet 391, 2128–2139 (2018).

16. Jorissen, R. N., Sakthianandeswaren, A. & Sieber, O. M. Immunoscore—has it scored for colon cancer precision medicine? Ann. Transl. Med. 6, S23 (2018).

17. Gough, A. et al. High-content analysis with cellular and tissue systems biology:a bridge between cancer cell biology and tissue-based diagnostics. in The Molecular Basis of Cancer 369–392 (Elsevier, 2015).

18. Schubert, W. et al. Analyzing proteome topology and function by automated multidimensional fluorescence microscopy. Nat. Biotechnol. 24, 1270 (2006).

19. Lin, J.-R., Fallahi-Sichani, M. & Sorger, P. K. Highly multiplexed imaging of single cells using a high-throughput cyclic immunofluorescence method. Nat. Commun. 6, 8390 (2015).

20. Gerdes, M. J. et al. Highly multiplexed single-cell analysis of formalin-fixed, paraffin-embedded cancer tissue. Proc. Natl. Acad. Sci. U. S. A. 110, 11982–11987 (2013).

21. Goltsev, Y. et al. Deep Profiling of Mouse Splenic Architecture with CODEX Multiplexed Imaging. Cell 174, 968–981.e15 (2018).

22. Buchberger, A. R., DeLaney, K., Johnson, J. & Li, L. Mass Spectrometry Imaging: A Review of Emerging Advancements and Future Insights. Anal. Chem. 90, 240–265 (2018).

23. Chen, K. H., Boettiger, A. N., Moffitt, J. R., Wang, S. & Zhuang, X. Spatially resolved, highly multiplexed RNA profiling in single cells. Science (80-.). 348, aaa6090 (2015).

24. Wang, G., Moffitt, J. R. & Zhuang, X. Multiplexed imaging of high-density libraries of RNAs with MERFISH and expansion microscopy. Sci. Rep. 8, 4847 (2018).

25. Lubeck, E., Coskun, A. F., Zhiyentayev, T., Ahmad, M. & Cai, L. Single-cell in situ RNA profiling by sequential hybridization. Nat. Methods 11, 360–361 (2014).

26. Bubendorf, L., Nocito, A., Moch, H. & Sauter, G. Tissue microarray (TMA) technology: miniaturized pathology archives for high-throughput in situ studies. J. Pathol. 195, 72–79 (2001).

27. Kendall, M. G. A new measure of rank correlation. Biometrika 30, 81–93 (1938).

28. Chambers, J. M. & Hastie, T. J. Statistical Models. in Statistical Models in S 13–44 (1992).

29. Wang, X. et al. Hypoxic Tumor-Derived Exosomal miR-301a Mediates M2 Macrophage Polarization via PTEN/PI3Kγ to Promote Pancreatic Cancer Metastasis. Cancer Res. 78, 4586 LP – 4598 (2018).

30. Maia, J., Caja, S., Strano Moraes, M. C., Couto, N. & Costa-Silva, B. Exosome-Based Cell-Cell Communication in the Tumor Microenvironment. Frontiers in Cell and Developmental Biology 6, 18 (2018).

31. Goeman, J. J. L1 Penalized Estimation in the Cox Proportional Hazards Model. Biometrical J. 52, 70–84 (2010).

32. Simon, N., Friedman, J., Hastie, T. & Tibshirani, R. Regularization Paths for Cox’s Proportional Hazards Model via Coordinate Descent. J. Stat. Softw. 39, 1–13 (2011).

33. Youden, W. J. Index for rating diagnostic tests. Cancer 3, 32–35 (1950).

34. Peddareddigari, V. G., Wang, D. & Dubois, R. N. The tumor microenvironment in colorectal carcinogenesis. Cancer Microenviron. 3, 149–166 (2010).

35. Therneau, T. M. & Grambsch, P. M. Modeling Survival Data: Extending the Cox Model. (Springer-Verlag, 2000).

36. Dunn, O. J. Multiple Comparisons Using Rank Sums. Technometrics 6, 241–252 (1964).

37. Benson, A. B., et al. Colon Cancer, Version 1.2017, NCCN Clinical Practice Guidelines in Oncology. J. Natl. Compr. Cancer Netw. J Natl Compr Canc Netw 15, 370–398 (2017).

38. Benson, A. B. et al. American Society of Clinical Oncology Recommendations on Adjuvant Chemotherapy for Stage II Colon Cancer. J. Clin. Oncol. 22, 3408–3419 (2004).

39. Group, on behalf of the E. G. W. et al. Early colon cancer: ESMO Clinical Practice Guidelines for diagnosis, treatment and follow-up†. Ann. Oncol. 24, vi64–vi72 (2013).

40. Varghese, A. Chemotherapy for Stage II Colon Cancer. Clin. Colon Rectal Surg. 28, 256–261 (2015).

41. Gan, S., Wilson, K. & Hollington, P. Surveillance of patients following surgery with curative intent for colorectal cancer. World J. Gastroenterol. 13, 3816–3823 (2007).

42. Ryuk, J. P. et al. Predictive factors and the prognosis of recurrence of colorectal cancer within 2 years after curative resection. Ann. Surg. Treat. Res. 86, 143–151 (2014).

43. Hammond, K. & Margolin, D. A. The role of postoperative surveillance in colorectal cancer. Clin. Colon Rectal Surg. 20, 249–254 (2007).

44. Heagerty, P. J., Lumley, T. & Pepe, M. S. Time-Dependent ROC Curves for Censored Survival Data and a Diagnostic Marker. Biometrics 56, 337–344 (2000).

45. Lin, J. Divergence measures based on the Shannon entropy. IEEE Trans. Inf. Theory 37, 145–151 (1991).

46. Sevinsky, C., et al. Abstract 1467: Multiplexed immunofluorescence quantitation and validation of multiple immune cell types in colon cancer epithelium and stroma. Cancer Res. 76, 1467 LP – 1467 (2016).

47. Tommelein, J. et al. Cancer-associated fibroblasts connect metastasis-promoting communication in colorectal cancer. Front. Oncol. 5, 63 (2015).

48. Bradley, C. A. et al. Transcriptional upregulation of c-MET is associated with invasion and tumor budding in colorectal cancer. Oncotarget 7, 78932–78945 (2016).

49. Szklarczyk, D. et al. STRING v11: protein-protein association networks with increased coverage, supporting functional discovery in genome-wide experimental datasets. Nucleic Acids Res. 47, D607–D613 (2019).

50. Kanehisa, M., Sato, Y., Furumichi, M., Morishima, K. & Tanabe, M. New approach for understanding genome variations in KEGG. Nucleic Acids Res. 47, D590–D595 (2019).

51. Colakoglu, T. et al. Clinicopathological significance of PTEN loss and the phosphoinositide 3-kinase/Akt pathway in sporadic colorectal neoplasms: is PTEN loss predictor of local recurrence? Am. J. Surg. 195, 719–725 (2008).

52. Vu, T. & Datta, P. K. Regulation of EMT in Colorectal Cancer: A Culprit in Metastasis. Cancers (Basel). 9, 171 (2017).

53. Bonnans, C., Chou, J. & Werb, Z. Remodelling the extracellular matrix in development and disease. Nat. Rev. Mol. Cell Biol. 15, 786–801 (2014).

54. Morgan, M. R., Humphries, M. J. & Bass, M. D. Synergistic control of cell adhesion by integrins and syndecans. Nat. Rev. Mol. Cell Biol. 8, 957–969 (2007).

55. Chen, C., Zhao, S., Karnad, A. & Freeman, J. W. The biology and role of CD44 in cancer progression: therapeutic implications. J. Hematol. Oncol. 11, 64 (2018).

56. Theocharis, A. D., Skandalis, S. S., Tzanakakis, G. N. & Karamanos, N. K. Proteoglycans in health and disease: novel roles for proteoglycans in malignancy and their pharmacological targeting. FEBS J. 277, 3904–3923 (2010).

57. Krashin, E., Piekiełko-Witkowska, A., Ellis, M. & Ashur-Fabian, O. Thyroid Hormones and Cancer: A Comprehensive Review of Preclinical and Clinical Studies. Frontiers in Endocrinology 10, 59 (2019).

58. Brown, A. R., Simmen, R. C. M. & Simmen, F. A. The role of thyroid hormone signaling in the prevention of digestive system cancers. Int. J. Mol. Sci. 14, 16240–16257 (2013).

59. Iftekhar, A., Sperlich, A., Janssen, K.-P. & Sigal, M. Microbiome and Diseases: Colorectal Cancer BT - The Gut Microbiome in Health and Disease. in (ed. Haller, D.) 231–249 (Springer International Publishing, 2018). doi:10.1007/978-3-319-90545-7_15

60. Dejea, C., Wick, E. & Sears, C. L. Bacterial oncogenesis in the colon. Future Microbiol. 8, 445–460 (2013).

61. Vaupel, P. The Role of Hypoxia-Induced Factors in Tumor Progression. Oncol. 9, 10–17 (2004).

62. Ross, J. S. et al. Targeting HER2 in colorectal cancer: The landscape of amplification and short variant mutations in ERBB2 and ERBB3. Cancer 124, 1358–1373 (2018).

63. Park, J. H. et al. Mismatch repair status in patients with primary operable colorectal cancer: associations with the local and systemic tumour environment. Br. J. Cancer 114, 562–570 (2016).

64. Chen, X. & Song, E. Turning foes to friends: targeting cancer-associated fibroblasts. Nat. Rev. Drug Discov. 18, 99–115 (2019).

65. Haviv, I., Polyak, K., Qiu, W., Hu, M. & Campbell, I. Origin of carcinoma associated fibroblasts. Cell Cycle 8, 589–595 (2009).

66. Vergadi, E., Ieronymaki, E., Lyroni, K., Vaporidi, K. & Tsatsanis, C. Akt Signaling Pathway in Macrophage Activation and M1/M2 Polarization. J. Immunol. 198, 1006 LP – 1014 (>2017).

67. Sveen, A., Kopetz, S. & Lothe, R. A. Biomarker-guided therapy for colorectal cancer: strength in complexity. Nat. Rev. Clin. Oncol. (2019).

68. Spagnolo, D. et al. Pointwise mutual information quantifies intratumor heterogeneity in tissue sections labeled with multiple fluorescent biomarkers. J. Pathol. Inform. 7, 47 (2016).

69. Janiszewska, M. et al. In situ single-cell analysis identifies heterogeneity for PIK3CA mutation and HER2 amplification in HER2-positive breast cancer. Nat. Genet. 47, 1212– 1219 (2015).

70. Balkwill, F. R., Capasso, M. & Hagemann, T. The tumor microenvironment at a glance. J. Cell Sci. 125, 5591 LP – 5596 (2012).

## References

1. Gerdes, M. J. et al. Highly multiplexed single-cell analysis of formalin-fixed, paraffin-embedded cancer tissue. Proc. Natl. Acad. Sci. U. S. A. 110, 11982–11987 (2013).

2. Bello, M., Can, A. & Tao, X. Accurate registration and failure detection in tissue micro array images. in 2008 5th IEEE International Symposium on Biomedical Imaging: From Nano to Macro 368–371 (2008). doi:10.1109/ISBI.2008.4541009

3. Woolfe, F., Gerdes, M., Bello, M., Tao, X. & Can, A. Autofluorescence Removal by Non-Negative Matrix Factorization. IEEE Trans. Image Process. 20, 1085–1093 (2011).

4. Pang, Z., Laplante, N. E. & Filkins, R. J. Dark pixel intensity determination and its applications in normalizing different exposure time and autofluorescence removal. J. Microsc. 246, 1–10 (2012).

5. Chambers, J. M. & Hastie, T. J. Statistical Models. in Statistical Models in S 13–44 (1992).

6. Simon, N., Friedman, J., Hastie, T. & Tibshirani, R. Regularization Paths for Cox’s Proportional Hazards Model via Coordinate Descent. J. Stat. Softw. 39, 1–13 (2011).

7. Friedman, J., Hastie, T. & Tibshirani, R. Regularization Paths for Generalized Linear Models via Coordinate Descent. J. Stat. Softw. 33, 1–22 (2010).

8. Harrell, F. E. Regression Modeling Strategies. (Springer, 2015).

9. Whittaker, J. Graphical models in applied multivariate statistics. (Wiley, 1990).

10. Szklarczyk, D. et al. The STRING database in 2017: quality-controlled protein-protein association networks, made broadly accessible. Nucleic Acids Res. 45, D362–D368 (2017).

11. Szklarczyk, D. et al. STRING v11: protein-protein association networks with increased coverage, supporting functional discovery in genome-wide experimental datasets. Nucleic Acids Res. 47, D607–D613 (2019).

## References

s1. Francipane, M. G. & Lagasse, E. mTOR pathway in colorectal cancer: an update. Oncotarget 5, 49– 66 (2014).

s2. Burotto, M., Chiou, V. L., Lee, J.-M. & Kohn, E. C. The MAPK pathway across different malignancies: a new perspective. Cancer 120, 3446–3456 (2014).

s3. Li, G.-M. Mechanisms and functions of DNA mismatch repair. Cell Res. 18, 85–98 (2008).

s4. Wu, C., Zhu, X., Liu, W., Ruan, T. & Tao, K. Hedgehog signaling pathway in colorectal cancer: function, mechanism, and therapy. Onco. Targets. Ther. 10, 3249–3259 (2017).

s5. Greenhough, A. et al. Cancer cell adaptation to hypoxia involves a HIF-GPRC5A-YAP axis. EMBO Mol. Med. 10, e8699 (2018).

s6. Watson, A. J. M. Apoptosis and colorectal cancer. Gut 53, 1701 LP – 1709 (2004).

s7. González-Trejo, S. et al. Baseline serum albumin and other common clinical markers are prognostic factors in colorectal carcinoma: A retrospective cohort study. Medicine (Baltimore). 96, (2017).

s8. Fujikawa, H. et al. Prognostic Impact of Preoperative Albumin–to–Globulin Ratio in Patients with Colon Cancer Undergoing Surgery with Curative Intent. Anticancer Res. 37, 1335–1342 (2017).

s9. Huang, H. et al. Validation of Prognosis Value of Cumulative Prognostic Scores Based on Serum High-Density Lipoprotein Cholesterol and Albumin Levels in Patients with Colorectal Cancer. J. Cancer 10, 35–42 (2019).

s10. Feng, W. et al. Role of glucose metabolism related gene GLUT1 in the occurrence and prognosis of colorectal cancer. Oncotarget 8, 56850–56857 (2017).

s11. Shen, Y.-M., Arbman, G., Olsson, B. & Sun, X.-F. Overexpression of GLUT1 in Colorectal Cancer is Independently Associated with Poor Prognosis. Int. J. Biol. Markers 26, 166–172 (2011).

s12. Ahopelto, K., Böckelman, C., Hagström, J., Koskensalo, S. & Haglund, C. Transketolase-like protein 1 expression predicts poor prognosis in colorectal cancer. Cancer Biol. Ther. 17, 163–168 (2016).

s13. Langbein, S. et al. Expression of transketolase TKTL1 predicts colon and urothelial cancer patient survival: Warburg effect reinterpreted. Br. J. Cancer 94, 578–585 (2006).

s14. Pan, S. et al. Decreased expression of ARHGAP15 promotes the development of colorectal cancer through PTEN/AKT/FOXO1 axis. Cell Death Dis. 9, 673 (2018).

s15. Farhan, M. et al. FOXO Signaling Pathways as Therapeutic Targets in Cancer. Int. J. Biol. Sci. 13, 815–827 (2017).

s16. Bullock, M. D. et al. FOXO3 expression during colorectal cancer progression: biomarker potential reflects a tumour suppressor role. Br. J. Cancer 109, 387–394 (2013).

s17. Li, X.-L., Zhou, J., Chen, Z.-R. & Chng, W.-J. P53 mutations in colorectal cancer - molecular pathogenesis and pharmacological reactivation. World J. Gastroenterol. 21, 84–93 (2015).

s18. Molinari, F. & Frattini, M. Functions and Regulation of the PTEN Gene in Colorectal Cancer. Front. Oncol. 3, 326 (2014).

s19. Ying, J. et al. WNT5A Exhibits Tumor-Suppressive Activity through Antagonizing the Wnt/β-Catenin Signaling, and Is Frequently Methylated in Colorectal Cancer. Clin. Cancer Res. 14, 55 LP – 61 (2008).

s20. Munz, M., Baeuerle, P. A. & Gires, O. The Emerging Role of EpCAM in Cancer and Stem Cell Signaling. Cancer Res. 69, 5627 LP – 5629 (2009).

s21. Greenhough, A. et al. The COX-2/PGE 2 pathway: key roles in the hallmarks of cancer and adaptation to the tumour microenvironment. Carcinogenesis 30, 377–386 (2009).

s22. Wang, D. & DuBois, R. N. The role of COX-2 in intestinal inflammation and colorectal cancer. Oncogene 29, 781–788 (2010).

s23. Lee, S. J. et al. c-MET Overexpression in Colorectal Cancer: A Poor Prognostic Factor for Survival. Clin. Colorectal Cancer 17, 165–169 (2018).

s24. Rasola, A. et al. A positive feedback loop between hepatocyte growth factor receptor and β-catenin sustains colorectal cancer cell invasive growth. Oncogene 26, 1078–1087 (2007).

s25. Shang, S., Hua, F. & Hu, Z.-W. The regulation of β-catenin activity and function in cancer: therapeutic opportunities. Oncotarget 8, 33972–33989 (2017).

s26. Gan, L. et al. Epigenetic regulation of cancer progression by EZH2: from biological insights to therapeutic potential. Biomark. Res. 6, 10 (2018).

s27. Vilorio-Marqués, L. et al. The role of EZH2 in overall survival of colorectal cancer: a meta-analysis. Sci. Rep. 7, 13806 (2017).

s28. Karantanos, T., Chistofides, A., Barhdan, K., Li, L. & Boussiotis, V. A. Regulation of T Cell Differentiation and Function by EZH2. Front. Immunol. 7, 172 (2016).

s29. Goswami, S. et al. Modulation of EZH2 expression in T cells improves efficacy of anti–CTLA-4 therapy. J. Clin. Invest. 128, 3813–3818 (2018).

s30. DuPage, M. et al. The chromatin-modifying enzyme Ezh2 is critical for the maintenance of regulatory T cell identity after activation. Immunity 42, 227–238 (2015).

s31. Simiczyjew, A., Mazur, A. J., Popow-Woźniak, A., Malicka-Błaszkiewicz, M. & Nowak, D. Effect of overexpression of β- and γ-actin isoforms on actin cytoskeleton organization and migration of human colon cancer cells. Histochem. Cell Biol. 142, 307–322 (2014).

s32. Dhawan, P. et al. Claudin-1 regulates cellular transformation and metastatic behavior in colon cancer. J. Clin. Invest. 115, 1765–1776 (2005).

s33. Kim, S. A. et al. Loss of CDH1 (E-cadherin) expression is associated with infiltrative tumour growth and lymph node metastasis. Br. J. Cancer 114, 199–206 (2016).

s34. Willis, N. D. et al. Lamin A/C is a risk biomarker in colorectal cancer. PLoS One 3, e2988–e2988 (2008).

s35. Belt, E. J. T. et al. Loss of lamin A/C expression in stage II and III colon cancer is associated with disease recurrence. Eur. J. Cancer 47, 1837–1845 (2011).

s36. Jain, R., Fischer, S., Serra, S. & Chetty, R. The Use of Cytokeratin 19 (CK19) Immunohistochemistry in Lesions of the Pancreas, Gastrointestinal Tract, and Liver. Appl. Immunohistochem. Mol. Morphol. 18, (2010).

s37. Clausen, M. V, Hilbers, F. & Poulsen, H. The Structure and Function of the Na,K-ATPase Isoforms in Health and Disease. Front. Physiol. 8, 371 (2017).

s38. Wang, J. P. & Hielscher, A. Fibronectin: How Its Aberrant Expression in Tumors May Improve Therapeutic Targeting. J. Cancer 8, 674–682 (2017).

s39. Tanjore, H. & Kalluri, R. The role of type IV collagen and basement membranes in cancer progression and metastasis. Am. J. Pathol. 168, 715–717 (2006).

s40. Ardito, F., Giuliani, M., Perrone, D., Troiano, G. & Lo Muzio, L. The crucial role of protein phosphorylation in cell signaling and its use as targeted therapy (Review). Int. J. Mol. Med. 40, 271–280 (2017).

s41. Hessman, C. J., Bubbers, E. J., Billingsley, K. G., Herzig, D. O. & Wong, M. H. Loss of expression of the cancer stem cell marker aldehyde dehydrogenase 1 correlates with advanced-stage colorectal cancer. Am. J. Surg. 203, 649–653 (2012).

s42. Douville, J., Beaulieu, R. & Balicki, D. ALDH1 as a Functional Marker of Cancer Stem and Progenitor Cells. Stem Cells Dev. 18, 17–26 (2008).

s43. Nelson, B. H. CD20^+^ B Cells: The Other Tumor-Infiltrating Lymphocytes. J. Immunol. 185, 4977 LP – 4982 (2010).

s44. Berntsson, J., Nodin, B., Eberhard, J., Micke, P. & Jirström, K. Prognostic impact of tumour-infiltrating B cells and plasma cells in colorectal cancer. Int. J. Cancer 139, 1129–1139 (2016).

s45. Pinto, M. L. et al. The Two Faces of Tumor-Associated Macrophages and Their Clinical Significance in Colorectal Cancer. Front. Immunol. 10, 1875 (2019).

s46. Lu, J. et al. Endothelial cells promote the colorectal cancer stem cell phenotype through a soluble form of Jagged-1. Cancer Cell 23, 171–185 (2013).

s47. Chu, P. G. & Arber, D. A. CD79: A Review. Appl. Immunohistochem. Mol. Morphol. 9, (2001).

s48. Cavalleri, T. et al. Combined Low Densities of FoxP3^+^ and CD3^+^ Tumor-Infiltrating Lymphocytes Identify Stage II Colorectal Cancer at High Risk of Progression. Cancer Immunol. Res. 7, 751 LP – 758 (2019).

s49. Ziai, J. et al. CD8+ T cell infiltration in breast and colon cancer: A histologic and statistical analysis. PLoS One 13, e0190158–e0190158 (2018).

s50. Zhang, S. et al. CCL5-deficiency enhances intratumoral infiltration of CD8+ T cells in colorectal cancer. Cell Death Dis. 9, 766 (2018).

s51. Schweizer, J. et al. New consensus nomenclature for mammalian keratins. J. Cell Biol. 174, 169– 174 (2006).

s52. Liu, T., Zhou, L., Li, D., Andl, T. & Zhang, Y. Cancer-Associated Fibroblasts Build and Secure the Tumor Microenvironment. Frontiers in Cell and Developmental Biology 7, 60 (2019).

s53. Nishishita, R. et al. Expression of cancer-associated fibroblast markers in advanced colorectal cancer. Oncol. Lett. 15, 6195–6202 (2018).

s54. Mi, L. et al. The metastatic suppressor NDRG1 inhibits EMT, migration and invasion through interaction and promotion of caveolin-1 ubiquitylation in human colorectal cancer cells. Oncogene 36, 4323–4335 (2017).

